# The Nonequilibrium Mechanism of Noise Enhancer synergizing with Activator in HIV Latency Reactivation

**DOI:** 10.1101/2020.01.14.905653

**Authors:** Xiaolu Guo, Tao Tang, Minxuan Duan, Lei Zhang, Hao Ge

## Abstract

Reactivating HIV latency and then simultaneously eliminating it by antiretroviral therapy has become a leading strategy in curing HIV. Recently, single-cell screening experiments have shown the drug synergy between two categories of biomolecules, Activators (AC) and Noise Enhancers (NE): NE can amplify the reactivation of latent HIV induced by AC, although NE itself cannot reactivate HIV latency. Based on an established LTR-two-state effective model, we uncover two necessary conditions for this type of drug synergy: The decreasing of the turning-on rate of LTR induced by NE is highly inhibited when presented with AC; The timescale of LTR turning off without AC/NE presented should be no slower than the timescale of Tat transactivation. Then we propose a detailed LTR-four-state mechanistic model with both AC/NE regulation kinetics and Tat transactivation circuit. We show that, in order to achieve drug synergy, the system of HIV gene state transition must operate out of thermodynamic equilibrium, which is caused by energy input. The direction of energy input determines whether the inhibition of NE upon the reactivation rate of LTR-off states (unbinding of RNAP) can be successfully prevented in the presence of AC. The drug synergy can also be significantly enhanced if the energy input is appropriately distributed to more than one reaction. Our model reveals a generic nonequilibrium mechanism underpinning the noise enhanced drug synergy, which may apply to identify the same drug synergy on reactivating a diverse class of lentivirus latency.

**Significance Statement:** The “kick and kill” strategy has become a promising way to cure HIV by eliminating latent HIV reservoirs, the main barrier to a clinical cure. Two categories of biomolecules, Activators (AC) and Noise Enhancers (NE), have been found to have synergy on reactivating latent HIV (kick strategy). We uncover the underlying non-equilibrium mechanism of such drug synergy by developing mathematical models based on genetic regulatory kinetics. We find that controlling the magnitude and direction of energy input into genetic regulatory kinetics can prevent NE from reducing the turn-on rate of the inactivated gene state in the presence of AC, which produces the synergy. This general nonequilibrium mechanism can be useful for identifying other drug synergies on lentivirus latency reactivation.

## Introduction

At the end of 2017, more than 36 million people were estimated to be infected with HIV (1). After HIV infects CD4+ cells, it can replicate or enter proviral latency (Figure 1A). Latent HIV reservoirs are the main obstacle to achieving a clinical cure (2). Reactivating latent HIV, quickly followed by antiretroviral therapy, has become a promising way to cure HIV-infected patients (3). Recently, synergistic combinations of noise enhancers and activators drugs were reported to beat other reactivation cocktails in reactivating HIV latency, yet induced less cytotoxicity (4). Two specific types of drugs used in (4): Activators (AC), a small biological molecule that increases the average expression level of HIV proteins, and Noise Enhancers (NE), a different type of molecule that increases the noise of HIV protein expression but does not affect the average expression level. While NE itself cannot reactivate latent HIV, it was shown to amplify AC-induced reactivation of HIV significantly (4). The synergy from adding NE with AC to latent HIV is shown in Figure 1B. However, the biomolecular mechanisms underlying the synergy between AC and NE have not been fully resolved.

**Figure 1.**
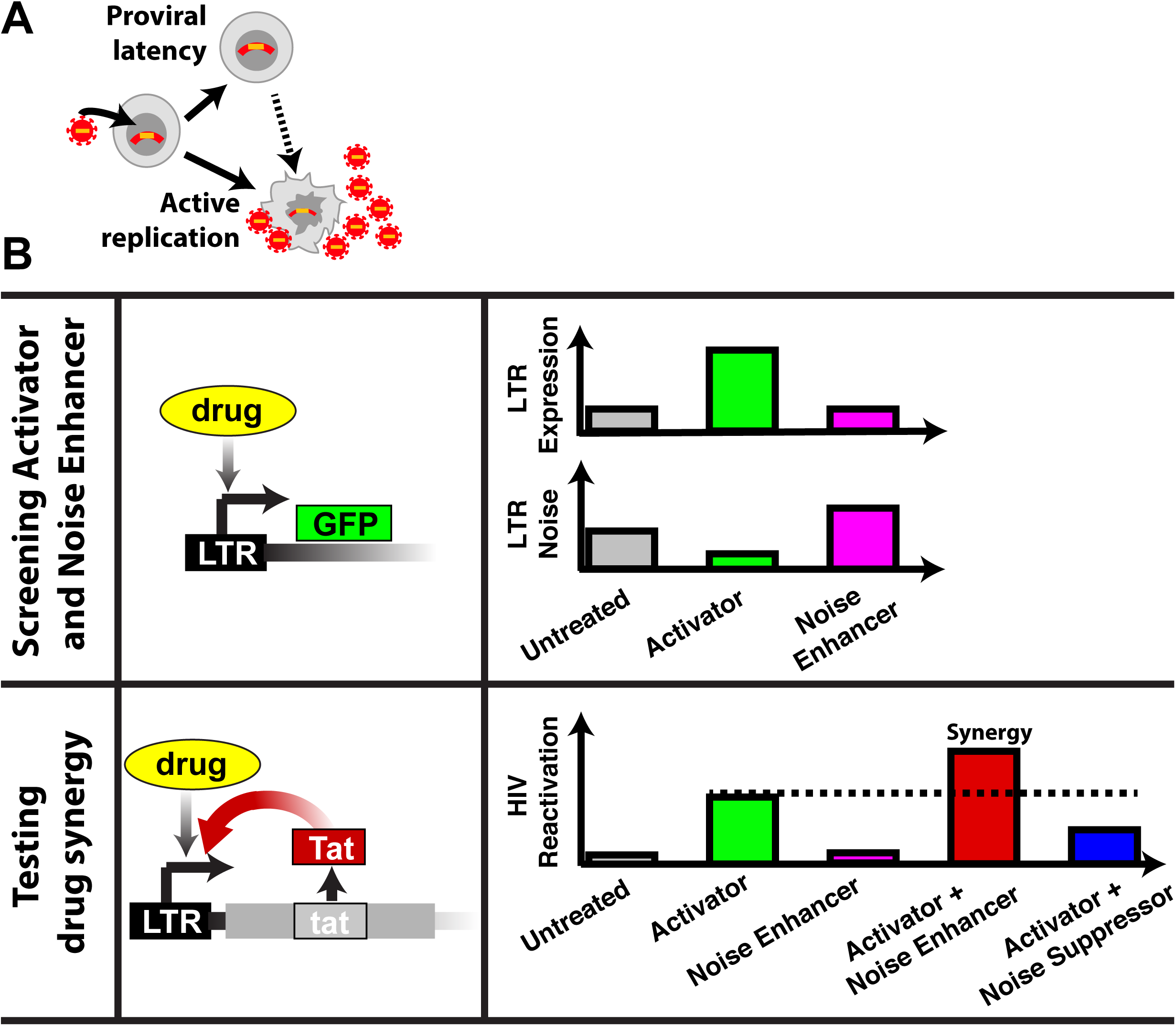
HIV-infected cell fates and biological function of biomolecules reactivating latent HIV. **(A)** Schematic of different fates of cells infected by HIV: HIV active replication, HIV proviral latency, and HIV latency reactivation (adapted from Figure 1A of (51)). **(B)** Diagram of experimentally screening Activator (AC) and Noise Enhancer (NE) (up) and experimental synergy test on the reactivation of latent HIV after adding AC or/and NE (down). In experiments, ACs and NEs were selected by to detecting the mean and noise of LTR expression using cells infected by the LTR-GFP vector. The synergy between ACs and NEs on HIV reactivation was tested using cells infected by full-length HIV with Tat transactivation. Untreated (grey bar) represents a control group. Adding Activator (green bar) increases LTR expression compared to the control group. Adding Noise Enhancer (magenta bar) increases LTR noise compared to the control group. Adding AC and NE simultaneously (red bar) has a strong synergy on HIV reactivation (increases Reactivation of latent HIV infected Cells). Adding AC and Noise Suppressors (NS, blue bar) has a depressing effect on HIV latency reactivation compared to adding AC only. (51)

The main ingredients of the HIV regulatory loop are the promoter long terminal repeats (LTR) and the Tat transactivation on LTR. LTR is the promoter of the HIV genome and has a larger expression noise than promoters of human genes (5, 6). Nucleosomes associated with the LTR often block the full transcription by RNA polymerase (RNAP), resulting in a low basal expression rate. The rarely produced Tat protein complexes with CDK9 and CyclinT1 to form the *positive transcriptional elongation factor b* (pTEFb). pTEFb can bind to the transactivation response element (TAR) on the initially transcribed part of the HIV mRNA and remodel downstream nucleosomes. This remodeling assists the elongation of the mRNA, thus forming positive feedback (7-9). In addition, a bimodal gene expression (“phenotypic bifurcation” (10)) pattern was found in the offspring of defective-HIV infected cells with initially intermediate expression (10). However, it was reported that the cooperativity coefficient (Hill coefficient) of Tat is only one (11), which means that the mean-field deterministic dynamics of HIV gene expression is monostable. This is distinct from the genetic toggle switch of the lambda phage regulatory loop with stronger feedback and bi-stable deterministic dynamics (12), or an oscillatory network with negative feedback and a limit cycle (13). The deterministic dynamics of the HIV regulatory network is insufficient to explain the observed bimodality, and a stochastic description may be required. By combining the bimodality and noisiness of HIV promoter gene expression, Weinberger et al. found that the bimodality arises from a very slow rate of switching on LTR expression (10), resulting in a much noisier dynamics than those observed for normal human promoters. On the other hand, functions of certain Activators and Noise Enhancers are partially known. For example, as Activators, TNF and prostratin can activate the transcriptional factor NF-κB, and therefore antagonize HIV latency(14-16). Some of these Noise Enhancers, such as ethinyl estradiol, can influence HIV expression through another transcriptional factor SP1 or the structural state of chromatin (17, 18). The molecular mechanisms of most noise enhancers are still unclear, indicating the complicated regulation mechanism of HIV dynamics. However, no matter how complex it is, what these activators or noise enhancers regulated are just the rates between different gene/promoter states described by TF binding or structural difference. Therefore, the functions of Activators and Noise Enhancers can be analyzed by certain minimal but general models.

Thermodynamic energy dissipation plays a crucial role in bioactivities and bioreactions. General model considering the binding of multiple transcription factors (TF) under thermodynamic equilibrium in prokaryotic cells and the function that different pair TF interactions can achieve in gene expression of cells was already studied extensively (19, 20). However, in studies of eukaryotic transcriptional dynamics, a non-equilibrium mechanism is found necessary(21), and many far-from-equilibrium models have been proposed (22-25). In addition to biomolecule synthesis and cell motility, the regulatory function of a living cell, such as adaptation and precise control of oscillations was also found highly dissipative (26, 27). Hence, we are very curious about whether certain energy input is necessary for this type of drug synergy.

In addition, in a self-positive-feedback gene regulatory network, the timescale of DNA state transition, mRNA transcription, mRNA decay, protein translation, and protein decay, will influence the cell fate landscape and phenotype transitions (28-30). Post-integration HIV gene expression is one example system of the TF regulatory mechanism of gene expression with self-positive-feedback. Hence, we are also interested in how the timescales of gene-state transition and protein dynamics influence the drug synergy.

In this essay, we first investigate an established LTR-two-state model and find two necessary conditions for the synergy between AC and NE on reactivating latently infected HIV: (i) AC inhibits NE’s function of reducing the transition rate *k*_on_ for the gene-state reactivation; (ii) the rate *k*_off_ for the promoter to turn off is not slower than the rate for Tat transactivation when the promoter is already turned on. We then propose an LTR-four-state model and prove that the noise enhanced drug synergy is indeed a non-equilibrium phenomenon of the regulation of HIV promoter LTR. We show that controlling the magnitude and direction of the system energy input can deter NE from reducing the turned-on rate of inactivated gene state in the presence of AC, which induces the synergy between AC and NE on the LTR transcription. The drug synergy can be significantly enhanced when we distribute the total energy input into two specific different reactions. Moreover, we show that only when the timescale of DNA unbinding RNAP (*k*_off_ or *k*_unbindp_) is comparable to or faster than the timescale of Tat transactivation at gene activated state, the synergy of AC and NE on the transcription on genetic level can pass to the translation on protein level in the Tat self-positive-feedback HIV system.

## Materials and methods

To investigate the synergy between AC and NE, we first employ a well-established LTR-two-state model with Tat transactivation from the previous study (Figure 2A) (4, 31). In this model, LTR (the promoter of HIV) has two states: on state and off state. LTR can transit to the transcriptionally inactive state LTR-off at rate *k*_off_ and transit to the transcriptionally active state LTR-on at rate *k*_on_. Tat can transactivate LTR-on state promoting a higher transcription rate *k*_transact_ than the basal transcription rate *k*_m_ due to Tat’s enhancing LTR transcriptional elongation (See Supplementary Section 1.1 for more details). The values of parameters (*k*_on_, *k*_off_, *k*_transact_) are the same as those in (31), which is quantified by single-cell analysis (32-34).

**Figure 2.**
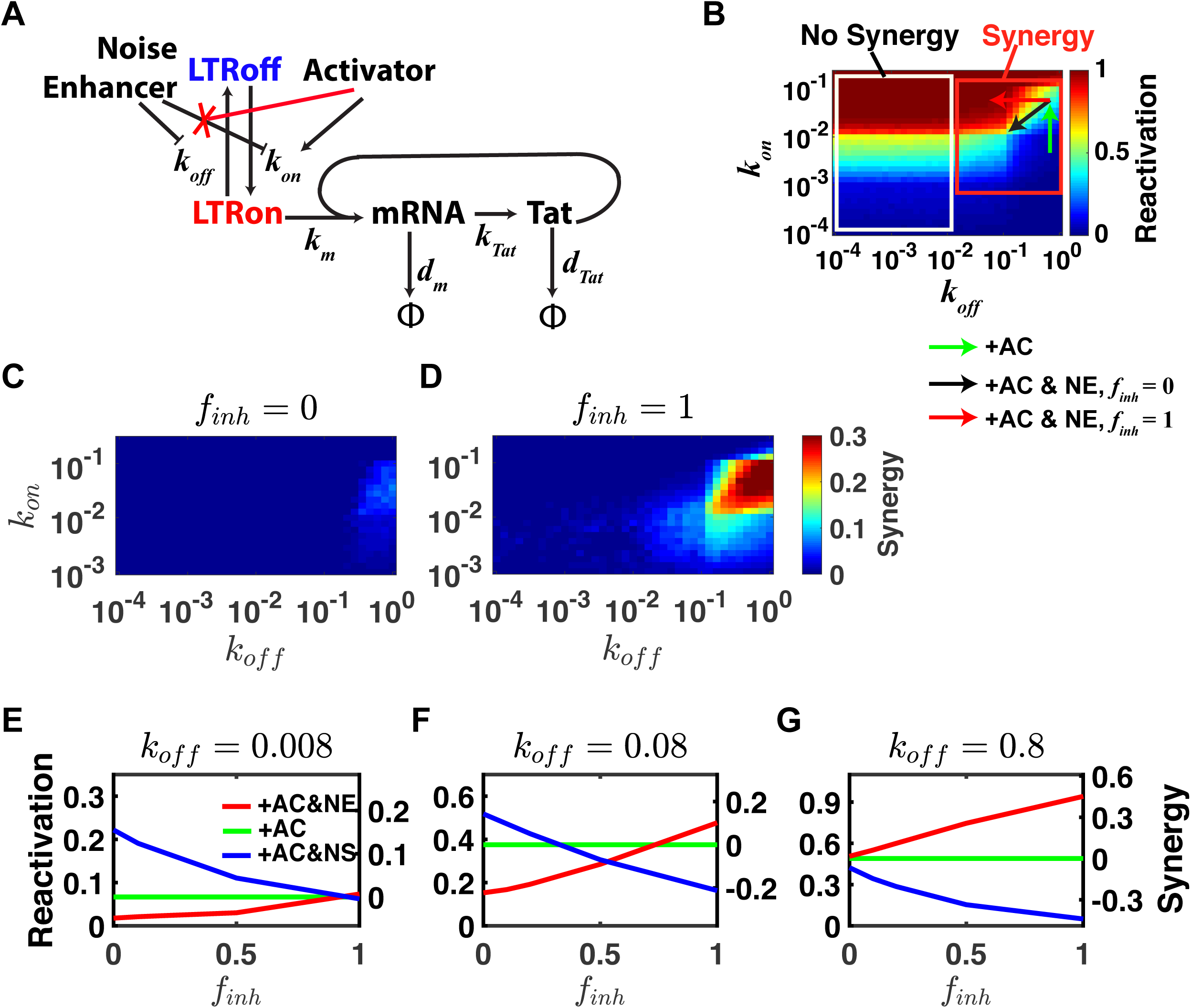
Two necessary conditions for drug synergy. **(A)** Modified from Figure 3A of (31). The LTR-two-state model with Tat feedback to explain the effects of NE and AC molecules on HIV. LTR has two states, on and off, which convert to each other at the rate of *k*_on_ and *k*_off_; the LTR-on state transcribes HIV mRNA at a rate *k*_m_, mRNA degrades at *d*_m_ rate or translates to protein at a rate of *k*_Tat_, and Tat degrades at rate *d*_Tat_. NE decreases *k*_on_ and *k*_off_ with their ratio fixed; AC molecules will increase *k*_on_; AC inhibits NE’s function on *k*_on_ when added together as assumed in (51). Tat transactivate LTR through enhancing the transcriptional rate *k*_trasact_. **(B)** The heat map of Reactivation across different values of LTR turning off rate *k*_off_ and LTR turning on rate *k*_on_. The green arrow corresponds to adding AC. Red arrow corresponds to adding NE only decreasing *k*_off_ without changing *k*_*on*_, i.e., AC inhibits NE’s function on *k*_on_. Black arrow corresponds to adding NE decreasing *k*_off_ and *k*_on_ with their ratio fixed, i.e., AC does not inhibit NE decreasing *k*_on_. (See Table S1-1 for parameter values) **(C-D)** The heat map of Synergy on Reactivation without AC’s inhibition on NE (*f*_inh_ = 0) and with AC’s complete inhibition on NE (*f*_inh_ = 1), respectively, across different values of *k*_off_ and *k*_on_ with AC added. The synergy is the reactivation with NE and AC added subtracting the reactivation with only AC added. (See Table S1-1 for parameter values) **(E-G)** The plots of synergy on reactivation changing with different *f*_*inh*_ values for small *k*_off_, intermediate *k*_off_, and large *k*_off_, respectively. Red lines indicate adding NE and AC simultaneously; green lines indicate adding AC only; blue lines indicate adding NS and AC simultaneously. (See Table S1-1 and Table S1-2 for parameter values)

As assumed in (4), adding only AC to the system increases *k*_on_, while adding only NE reduces *k*_on_ and *k*_off_ simultaneously with their ratio fixed. We use parameter *f*_inh_ to quantify the degree of AC’s inhibition upon the NE-induced reduction of *k*_on_, which is defined as

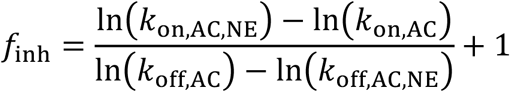

(See Supplementary Section 1.2-1.3 for more details). *f*_inh_ = 0 represents AC does not inhibit NE’s function of reducing *k*_on_ (Figure S1C, left panel), and *f*_inh_ > 0 means that AC inhibits NE’s function of reducing *k*_on_. Particularly, *f*_inh_ = 1 means that NE’s function of reducing *k*_on_ is fully inhibited by AC (Figure S1C, right panel).

We propose a more detailed LTR-four-state model with both transcript activation and Tat transactivation (Figure 3A, Figure S2A). As the HIV promoter, LTR controls HIV protein expression after the HIV genome integrates into the DNA of human CD4+ cells (35, 36). In this model, there are four different promoter states: LTR state is the free state; LTR* is the activated state but without RNAP binding; LTR-P and LTR*-P are the corresponding RNAP-bounded states. AC is assumed to promote LTR transiting to the activated state LTR*, e.g., LTR bound with NF-κB, and LTR* recruits RNA polymerase much easier than LTR itself, e.g., the NF-κB bound to LTR acting as a Transcription Factor (TF) to recruit RNA polymerase (RNAP) to LTR (37). Dar, Hosmane, Arkin, Siliciano and Weinberger (4) show that the screened Noise Enhancers have no effect on post transcription. Some NEs can increase transcription factors in cells, such as SP1 (17, 18). Similar to (4), we assume that NE can slow down the switching rates between LTR and LTR-P.

**Figure 3.**
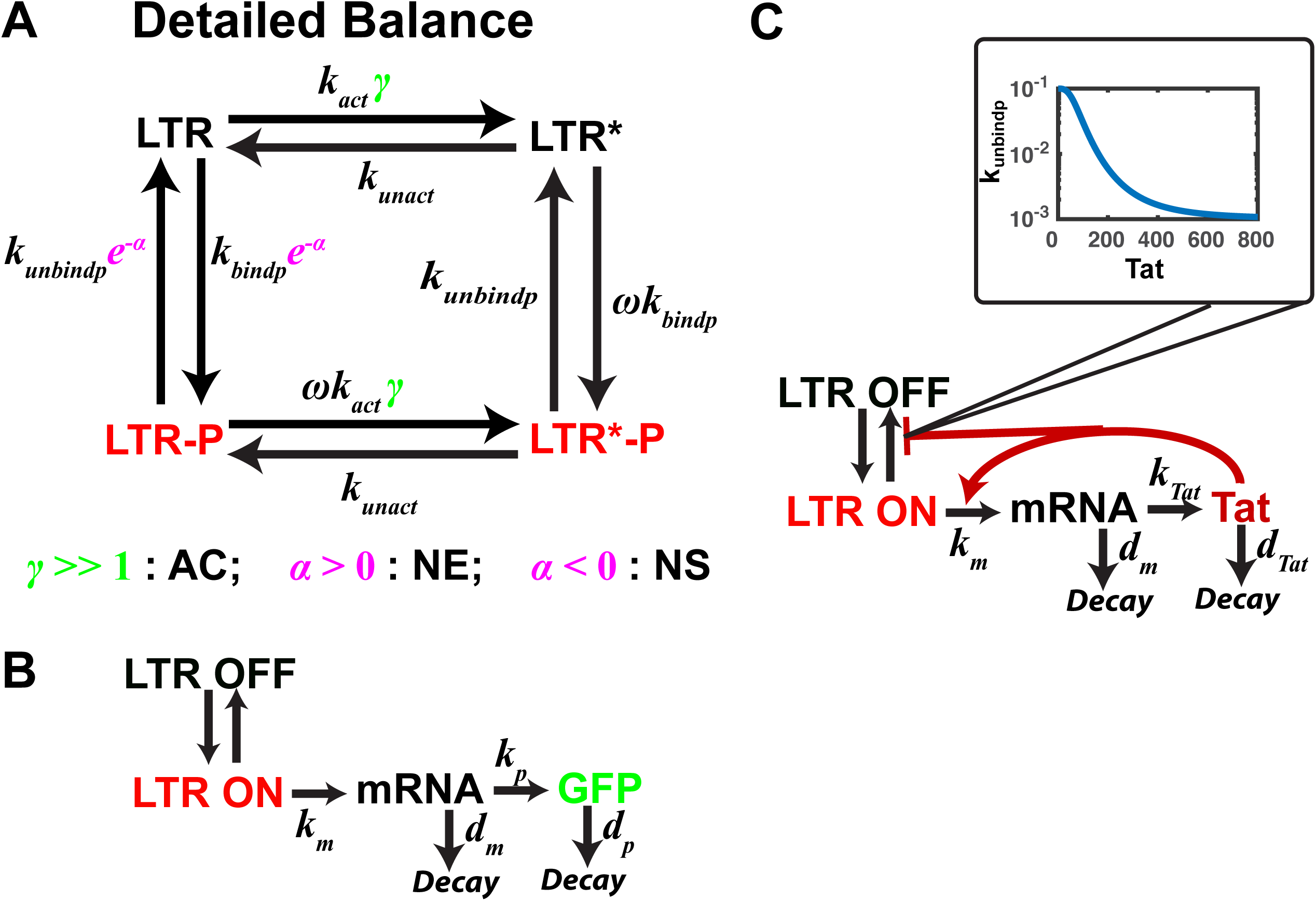
The LTR-four-state model under the detailed balance condition and transcription/translation modules without/with Tat transactivation. **(A)** Schematic of the LTR-four-state model with the Tat-feedback circuit. The LTR promoter is modeled to have four states: a transcriptional silence state (LTR state) extremely slowly binding RNAP polymerase or activation transcription factors such as NF-κB, an activated state (LTR*) such as LTR with NF-κB bond, a transcription-permissive state (LTR-P), a transcription-permissive state with NF-κB bond (LTR*-P). *k*_act_ is the rate for LTR binding NF-κB. *k*_unact_ is the rate for LTR unbinding NF-κB. *γ* models AC increases the rate for LTR binding NF-κB. *γ* = 1 for untreated HIV infected cells, and *γ* ≫ 1 for adding AC. *ω* is the attraction coefficient between NF-κB and RNAP, *ω* = 10 (*ω* > 1 means NF-κB attracts RNAP); *k*_bindp_ is the rate at which RNAP binds to LTR; *k*_unbindp_ is the rate at which RNAP unbinds from LTR; *α* is the noise attenuation factor (*α* > 0 corresponds to Noise Enhancer, and *α* < 0 corresponds to Noise Suppressor). The parameter setting here is under the detailed balance condition. The case of breaking the detailed balance is in Figure 5 and Table S2-1. **(B)** Schematic of the LTR-four-state model coupled with the transcription and translation module without feedback. LTR-on states (red, including LTR-P state and LTR*-P state) transcribes mRNA at rate *k*_m_; mRNA decays at rate *d*_m_ or can be translated at rate *k*_p_ into GFP; GFP decays at rate *d*_p_. **(C)** Schematic of the LTR-four-state model coupled with the transcription and translation module with the Tat-transactivation circuit. LTR-on states (red, including LTR-P state and LTR*-P state) transcribes mRNA at rate *k*_m_; mRNA decays at rate *d*_m_ or can be translated at rate *k*_Tat_ into Tat; Tat decays at rate *d*_Tat_; Tat has positive feedback on *k*_m_; Tat stabilizes the state of LTR-on state through negative feedback on *k*_unbindp_.

We use the Markov jumping process to model the transition among LTR states with AC and/or NE added (Figure 3A). The four states can mutually transit: in the absence of AC, RNAP binds to LTR at a relatively slow rate *k*_bindp_[*P*] and unbinds at a relatively fast rate *k*_unbindp_; LTR transit to LTR* state at an extremely slow rate *k*_act_ without AC, but at a much higher rate *k*_act_*γ* (*γ* ≫ 1) with AC presented; RNAP is recruited to LTR* at a higher rate *ωk*_bindp_, where *ω* is the attraction coefficient (*ω* > 1); when NE (or Noise Suppressor, NS) is added, LTR will bind and unbind RNAP at slower (faster for NS) rates (*k*_bindp_*e*^−*α*^, *k*_unbindp_*e*^−*α*^) with noise attenuation factor *α*. Note that the setup of the model showed in Figure 3A is under the assumption of the detailed balance. We will analyze both the models with or without this assumption in the following sections.

We also combine the LTR-four-state model with the transcription/translation module without Tat transactivation to characterize the procedure of screening AC and NE (Figure 3B). In screening experiments of AC and NE, the LTR-GFP vectors without Tat transactivation were used (4). For the LTR-GFP vectors, with RNAP bond to LTR (i.e., LTR-P state or LTR*-P state), the downstream DNA of LTR can be transcribed into mRNA at rate *k*_m_, independent of the GFP copy number. Then the mRNA can be translated into protein at rate *k*_GFP_. Also, mRNA and GFP will degrade at rate *d*_m_ and *d*_GFP_, respectively (Figure 3B).

In the experiments of testing the synergy between AC and NE on HIV latency reactivation, the full-length HIV with Tat transactivation was used to infect host cells instead of the LTR-GFP vector (4). For the full-length HIV vector, the Tat protein expressed by the HIV genome enhances transcriptional elongation by mediating RNAP hyperphosphorylation and thus transactivates the LTR promoter (38-40).

Based on the biological evidence and previous models(31, 41), we construct the Tat transactivation module (Figure 3C; see Supplementary Section 2.4 for details). Only at LTR-on states (LTR-P and LTR*-P), HIV genome can be transcribed into mRNA, and Tat transactivates LTR-on states in a Hill equation form:

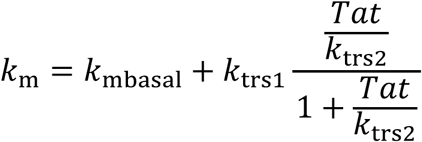

In addition, previous experiments also certified that Tat transactivation could stabilize HIV activation state autonomous from the host cellular state (31, 34). Based on this, we assume that Tat reduces the dissociating rate of RNAP from LTR, *k*_*unbindp*_, thereby stabilizing the HIV activation state (Figure 3C inset) (See Supplementary Section 2.4 and 2.12 for details and parameter value)

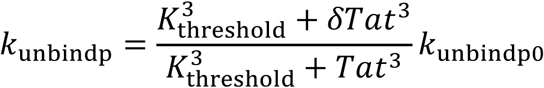

Then in the following sections, we use the LTR-four-state model coupled with Tat transactivation module to investigate how LTR expression and HIV fates are regulated by NE and AC under the Tat transactivation. We also simulate HIV expression trajectories in host T-cell from HIV latent initial state and calculate the percentage of cells expressing HIV protein reached a threshold (#Tat = 75) after a certain time (100 hours) as the reactivation ratio (Figure S2C; see Supplementary Section 2.7 for more details).

## Results

### The LTR-two-state model with significant large *k*_*off*_ can exhibit drug synergy between NE and AC

We simulate the LTR-two-state model showed in Figure 2A using Stochastic Simulation Algorithm (SSA) (42), calculating the reactivation of HIV latently infected T cells after adding NE and/or AC, with the above assumptions on AC’s and NE’s functions.

Through simulation, we find that AC’s inhibiting NE’s function on *k*_on_ is necessary for the synergy between AC and NE (Figure 2C-D). When AC has no inhibiting effect on NE, corresponding to *f*_inh_ = 0, there is no synergy between AC and NE in the whole reasonable parameter space (Figure 2C, Figure 2B black arrow). Only when AC inhibits NE’s function of reducing *k*_on_, there can be synergy between AC and NE on reactivating latent HIV (Figure 2D, Figure 2B red arrow).

We also find out that another necessary condition for NE having synergy with AC is *k*_off_ being greater than 10^−2^ hour^−1^, which is larger than the parameters used in previous studies (31). However, other important results shown in previous literature, the bimodal distribution of phenotype bifurcation (10) and HIV latency establishment operating autonomously from the host cellular state (31), are not sensitive to the increasing of *k*_off_ (Figure S1G-I). When *k*_off_ is no greater than 10^−2^, there is no synergy between AC and NE on reactivating latent HIV (Figure 2E; Figure S1E). However, if *k*_off_ is greater than 10^−2^, then NE can have synergy with AC on reactivating latent HIV (Figure 2F-G; Figure S1F). Furthermore, the synergy between AC and NE will increase with the inhibiting effect quantity *f*_inh_ > 0 only when *k*_off_ is sufficiently large, such as *k*_off_ = 0.8 > 10^−2^ (red line in Figure 2G). There will be no synergy between AC and NE with the inhibiting effect quantity *f*_inh_ > 0 if *k*_off_ < 10^−2^(red line in Figure 2E). Actually, in a latent HIV system, *k*_off_ is more likely to be larger than that previously used in literature. Direct sequencing of HIV integration sites showed that latent HIV frequently integrated into heterochromatin (38). The transcription is continuously turned off in heterochromatin, a tightly packed form of DNA (43). Thus, in latent HIV, LTR transcription-permissive state should be unstable, corresponding to larger *k*_off_, as one characteristic of the HIV integrating into heterochromatin.

In summary, we find two necessary conditions of drug synergy from our simulation results: (i) AC inhibits NE’s function of reducing *k*_on_; (ii) sufficiently large *k*_off_ (rate of LTR turning off or unbinding RNAP). However, this LTR-two-state model is oversimplified. We still do not know much about what causes AC inhibiting NE’s function of reducing *k*_on_ (the assumption made in (4)) and why NE synergizes with AC only if *k*_off_ is large.

### No drug synergy can be produced under the detailed balance condition in the LTR-four-state model coupling with Tat transactivation

Under the detailed balance condition (Figure 3A), our LTR-four-state model with the transcription/translation module without Tat transactivation (Figure 3B) illustrates that AC increases LTR mean expression level and that NE increases LTR expression noise (Figure S4C-D), which is consistent with the drug screening experimental results (Figure 1B) (4).

However, under the detailed balance condition, neither synergy between AC and NE nor depression of Noise Suppressor (NS) on AC reactivating latent HIV can be possible (Figure 4). This contradicts the experimental data showing that NEs enhance AC’s inducing latent HIV reactivation and that NSs reduce AC’s reactivating latent HIV (Figure 1B) (4). It is because under the detailed balance condition, the probability *P*_on_ of RNA polymerase binding to LTR (LTR-P and LTR*-P) (See Supplementary Section 2.1 and 2.5 for details) only depends on the equilibrium constants of each reaction, and NE or NS tunes the forward and backward rates simultaneously but keeps the equilibrium constant unvaried. This conclusion is not dependent on concrete models. It is a general physical and mathematical result. Hence it is not possible to build another even more complicated Detailed-Balanced model to overcome this obstacle.

**Figure 4.**
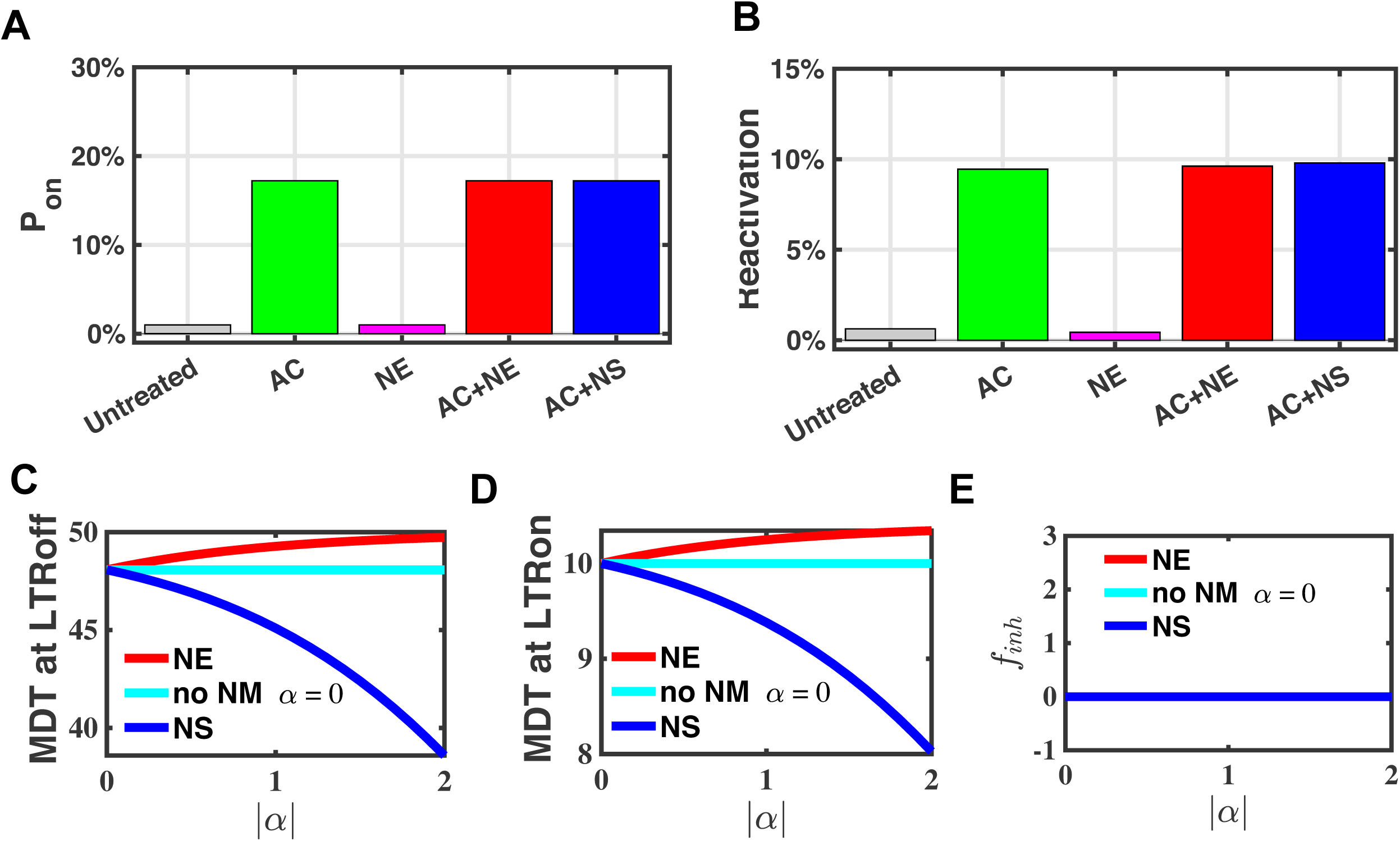
No synergy is predicted under the detailed balance condition. **(A-B)** Probability of LTR-on states (LTR-P state and LTR*-P state), *P*_on_, and Reactivation ratio of Latent HIV, respectively, in the Detailed-Balanced LTR-four-state model. Y-axis is the *P*_on_ and the reactivation ratio value, respectively, and X-axis is the categories of different combinations of ACs and NEs/NSs. Untreated (grey bars) corresponds to *γ* = 1, *α* = 0; adding only AC (green bars) corresponds to *γ* ≫ 1, *α* = 0; adding only NE (magenta bars) corresponds to *γ* = 1, *α* = 1; adding NE and AC (red bars) corresponds to *γ* ≫ 1, *α* = 1; adding NS and AC (blue bars) corresponds to *γ* ≫ 1, *α* = −1. (B) We use Stochastic Simulation Algorithm (SSA) to calculate the reactivation ratio of the LTR-four-state model coupled with Tat feedback. Reactivation Ratio is the ratio of reactivated HIV trajectory number by time 100h to all trajectory number, starting from latency state (LTR=1, all other species=0, simulated 5000 cells). **(C-D)** Mean Duration Time at LTR-off states and LTR-on states, respectively, under the detailed balance condition. **(E)** *f*_inh_, the degree of AC’s inhibition upon the reduction of *k*_on_ induced by NE under the detailed balance condition. We first calculate the reciprocal of the Mean Duration Time as the transition rate between LTR-on and LTR-off states, *λ*_on_ and *λ*_off_, respectively. Then we calculate using the formula: 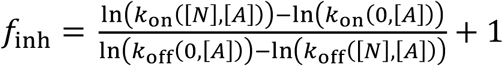. (C-E) The red lines correspond to adding AC and NE (*α* > 0); the cyan lines correspond to adding AC only (*α* = 0); the blue lines correspond to adding AC and NS (*α* < 0). (See Table S2-3 and Table S2-4 for parameter values.)

The above theory can be illustrated by our LTR-four-state model (Figure 3A, C, and Figure S2A, C) (See Supplementary Section 2.1,2.3 and 2.4 for details). Under the detailed balance condition, *P*_on_ keeps the same when both NE (or NS) and AC are added to the system (*γ* ≫ 1, *α* = 1 (or *α* = −1)) compared to when only AC is added (*γ* ≫ 1, *α* = 0) (Figure 4A; see Supplementary Section 2.5 for details). Thus, the Detailed-Balanced LTR-four-state model predicts no synergy between NE and AC and predicts that NS does not suppress the AC’s function of increasing *P*_on_ (Figure 4A), contradicting the experimentally observed synergy between AC and NE (Figure 1B) (4).

We couple the Detailed-Balanced LTR-four-state model with Tat transactivation, and there is still no synergy between Noise Enhancer and Activator illustrated by the reactivation ratio of latent HIV (Figure 4B; see Supplementary Section 2.7 for details). Note that the reactivation ratio of HIV is calculated dynamically during a finite time starting from the latent state, which is different from the steady-state probability *P*_on_. However, they are closely related to each other, since they both indicate the degree of reactivation for latent HIV.

In addition, we calculate the mean duration time (MDT) of both the LTR-off states (LTR and LTR*) and the LTR-on states (LTR-P and LTR*-P) (See Supplementary Section 2.8 for details). The reciprocals of the MDTs calculated from the LTR-four-state model can be regarded as the effective transition rates in the reduced LTR-two-state model with only the LTR-off and LTR-on state. Then we find out that AC can shorten the MDT at LTR-off states (Figure S4E, Figure S4F), and that NE can lengthen the MDT at both LTR-on states and LTR-off states with their ratio fixed (Figure 4C-D, S5G-H).

These results are consistent with the assumptions of the LTR-two-state model in the previous section. However, in this Detailed-Balanced model, the effective inhibiting effect quantity *f*_inh_ defined through the effective transition rates, always vanishes (Figure 4E; see Supplementary Section 2.9 for the exact definition of *f*_inh_). This confirms the LTR-two-state model predictions, that no synergy between AC and NE on reactivating latent HIV should be observed when *f*_inh_ = 0.

Hence, for Noise Enhancers synergize with ACs, the regulation of HIV gene expression must be a Non-Detailed-Balanced process with energy dissipation.

### The direction of the cycle flux caused by energy input in the Non-Detailed-Balanced LTR-four-state model determines the synergy

Inside a living cell, continuous energy consumption is necessary for executing different vital functions. We already know that systems with the drug synergy must be energy dissipative, but how energy input, i.e., breaking the detailed balance, influences the drug synergy remains poorly understood.

We mainly investigate how the cycle flux direction and the energy input distribution, as the features of the non-equilibrium system, effect the synergy. Breaking the detailed balance is equivalent to having non-vanishing cycle fluxes. In our LTR-four-state model, the cycle fluxes can be either counter-clockwise or clockwise. Energy input can be distributed on one or more reactions. Here, we first consider the case when energy input is only through one single reaction (Figure 5A; see Supplementary Section 2.2 and Table S2-1 for details). In the real biological system, the energy input can be realized through ATP hydrolysis or other reversible covalent modification (44).

**Figure 5.**
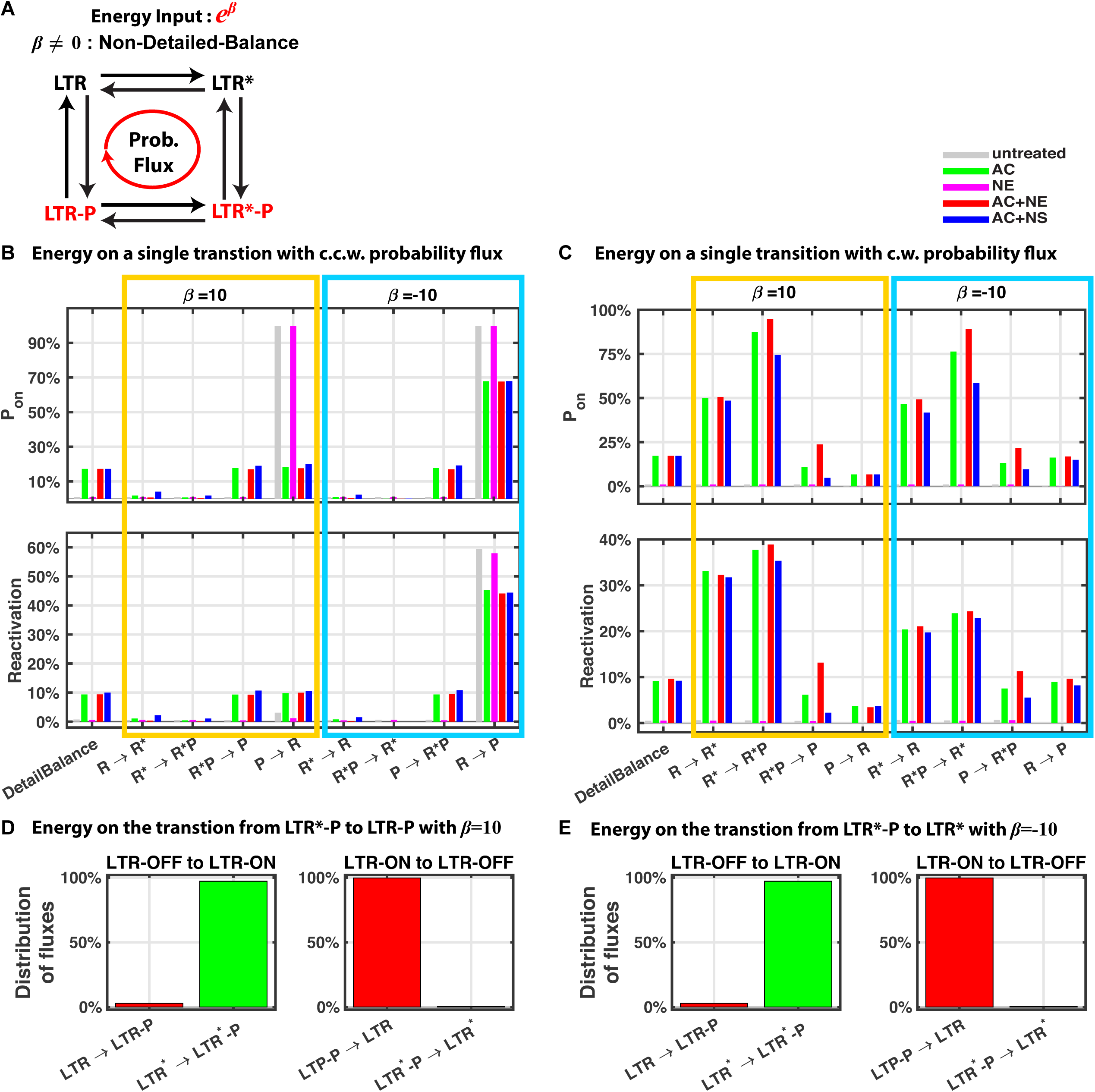
The LTR-four-state model with energy input on a single transition produces synergy between AC and NE if and only if the system has clockwise cyclic probability flux. **(A)** Schematic of the Non-Detailed-Balanced LTR-four-state model. Energy input can influence any single transition rate of the Detailed-Balanced LTR-four-state model with the corresponding rate multiplying by *e*^*β*^ or *e*^−*β*^. Such an energy input will cause clockwise (c.w.) cyclic probability flux or counter-clockwise (c.c.w.) cyclic probability flux. **(B-C)** Probability of LTR-on states, *P*_on_ (up panels), and Reactivation ratio of Latent HIV (down panels) calculated from the Non-Detailed-Balanced models with energy input on different single transitions causing a c.c.w. cyclic probability flux (B) or c.w. cyclic probability flux (C). R stands for LTR state; R* stands for LTR* state; P stands for LTR-P state; R*P stands for LTR*-P state. Each group of x-axis represents the Non-Detailed-Balanced model with the corresponding transition rate multiplying by *e*^*β*^ (*β* > 0 for orange groups, *β* < 0 for blue groups). (See Table S2-1 for model details. See Table S2-3 and Table S2-4 for parameter values.) **(D-E)** The distributions of fluxes for LTR turning on (left panels) and turning off (right panels). (See Table S2-1 and Supplementary Section 2.5-7 for model details. See Table S2-3 and Table S2-4 for parameter values.)

We prove that the Non-Detailed-Balanced LTR-four-state model can produce the drug synergy between NE and AC on *P*_on_, if and only if the direction of cycle flux is clockwise. Mathematical analysis (See Supplementary Section 2.10 for details) and numerical simulations illustrate the same phenomenon. The model with counter-clockwise cycle flux predicts no synergy between NE and AC on *P*_on_ or HIV latency reactivation, and no reduction of *P*_on_ or latent HIV reactivation is observed when NS is added with AC (Figure 5B). On the other hand, with clockwise cycle flux, the model predicts 100% of NE can synergize with AC on *P*_on_, and about 75% of NE synergize with AC on latent HIV reactivation (Figure 5C). 50% (4 out of 8) of the ways to break the detailed balance through a single reaction to produce a clockwise cyclic probability flux predicts significant synergy (> 5%) on *P*_on_ between NE and AC (Figure 5C up panel), and that NS reduces *P*_on_ with AC added. 25% (2 out of 8, increasing transition rate from LTR*-P to LTR-P or reducing transition rate from LTR-P to LTR*-P) of the ways predicts significant synergy (> 5%) on reactivation of latent HIV between NE and AC (Figure 5C down panel), and that NS reduces AC-induced HIV latency reactivation. In the experiments, around 80% of the NSs reduce AC-induced HIV latency reactivation, while more than 64% of NEs have synergistic effects with AC on reactivating latent HIV (4). Hence the results of our model are consistent with the experimental fact that the majority of NEs amplify AC reactivating latent HIV, while the majority of NSs suppresses reactivation of latent HIV with AC added. Thus, the Non-Detailed-Balanced LTR-four-state model reveals a general mechanism of the synergy between NEs and ACs on the reactivation of latent HIV, instead of a particular mechanism of a specific NE.

We also show that in the above cases producing significant drug synergy between AC and NE, the clockwise cyclic probability flux always promotes LTR turn on mainly through the LTR*-to-LTR*-P pathway strengthened by AC and turn off through the LTR-P-to-LTR pathway weakened directly by NE (Figure 5D-E). It explains why NE can further amply the HIV latency reactivation induced by AC, as long as the energy input provides clockwise cyclic probability flux.

In addition, for the equilibrium system, the distribution of the dwell time at LTR-off states is predicted to be monotonically decreasing and convex (45) (Figure S10, solid black lines). The monotonicity or convexity can be maintained for the non-equilibrium system with a low magnitude of energy input (small disturbance from the equilibrium system) (Figure S10, dashed red line). However, as the magnitude of energy dissipation increases, the nonmonotonicity or concavity of the distribution of dwell time could appear, similar to the phenomenon of phase transition (Figure S10D, H, dotted red line, and solid red line).

### The LTR-four-state model with distributed energy input may achieve much stronger synergy than that with energy input only on a single reaction

One possible strategy, through which strong synergy can be achieved, is to drive the LTR promoter to turn on mostly through the LTR-to-LTR*-to-LTR*-P pathway, whose rate can be significantly increased by AC, and to turn off mostly through the LTR*P-to-LTR-P-to-LTR pathway (Figure 5D-E), whose rate can be distinctly decreased by NE. This way, the promoter is more likely to transit to the state LTR-P rather than the state LTR* once it is at the state LTR*-P. Here, we build an EITST (Energy Input on the Two Specific Transition rates) LTR-four-state model, in which part of the energy input reduces the transition rate from LTR*-P to LTR* (*β*_2_) and the other part increases the transition rate from LTR*-P to LTR-P (*β*_1_) (Figure 6A; see Supplementary Section 2.2 for details), with the total energy fixed (*β*_1_ + *β*_2_ = *β*).

**Figure 6.**
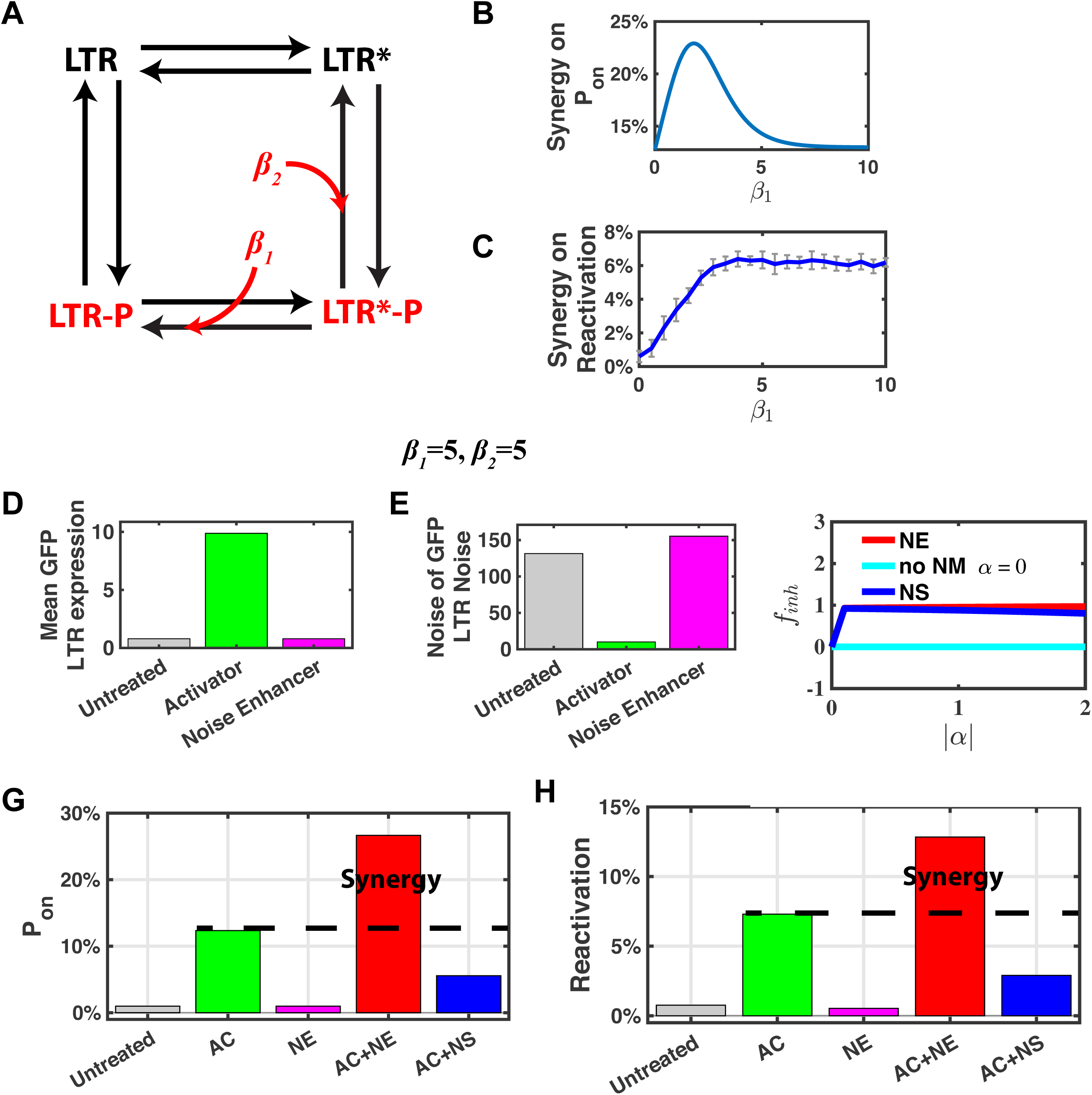
The LTR-four-state model with distributed energy input exhibit strong synergy between AC and NE. **(A)** Schematic of the EITST model with distributed energy input (*β* = *β*_1_ + *β*_2_ = 10). The first part of the energy *β*_1_ increases the LTR*-P-to-LTR-P transition rate by multiplying 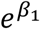, the other part of the energy *β*_2_ reduces the LTR*-P-to-LTR* transition rate by multiplying 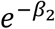. **(B-C)** The synergy on *P*_on_ (B) and HIV latency reactivation (C) vary with energy distribution (*β*_1_ + *β*_2_ = *β* = 10). (C) Each point is average of 250 simulation experimental data points with 10000 cells simulated for each simulation experiment. Error bars show the Standard Deviation. **(D-J)** *β*_1_ = *β*_2_ = 5. **(D-E)** The mean and noise of GFP expression calculated from the EITST model without positive feedback. **(F)** *f*_inh_, the degree of AC’s inhibition upon the reduction of *k*_on_ induced by NE of the EITST model. **(G-H)** Probability of LTR-on states (LTR-P state and LTR*-P state), *P*_on_, and Reactivation ratio of Latent HIV, calculated from the EITST model. (See Table S2-1 for model details. See Table S2-3, S2-4, and S2-5 for parameter values.)

We find that there is an optimal energy input distribution (*β*_1_ = 1.8, *β*_2_ = 8.2) for the system to perform the strongest synergy between AC and NE on *P*_on_ (Figure 6B). The drug synergy on HIV latency reactivation dependent on the energy input distribution is qualitatively quite similar. (Figure 6C). Overall, the certain distributed energy input with 0 < *β*_1_ < 10 may achieve stronger synergy than that on only a single reaction (*β*_1_ = 0 or *β*_1_ = 10). Without loss of generality, we set *β*_1_ = 5, *β*_2_ = 5 for the EITST LTR-four-state model, and all the following simulation results are based on this value.

In such a Non-Detailed-Balanced model, simulation results of adding AC or NE alone with GFP present are consistent with the drug screening experimental data, that AC increases LTR mean expression level and NE increases LTR expression noise (Figure 6D-E, Figure 1B).

The synergy between Noise Enhancer and Activator on both *P*_on_ (Figure 6G) and HIV latency reactivation (Figure 6H) are observed and much stronger than the scenario where energy input is only on a single reaction. Noise Enhancer can increase the HIV latency reactivation from approximately 7% when Activator is already added to 13% when both are added (Figure 6H). These numbers are quite similar to the best cases observed in experiments with prostratin as AC (experimental data from Figure 3A in (4)). And Noise suppressor reduces the AC-induced HIV latency reactivation from 7% to less than 1% (Figure 6H). This synergy between AC and NE on both *P*_on_ and HIV latency reactivation is found to be positively correlated with the magnitude |*β*| of the energy input, but will reach maximum (for *P*_on_) and saturation (for HIV latency reactivation) when |*β*| is getting sufficiently large (*β* > 5) (Figure S6A-B). In addition, the synergy is found to be positively correlated with the noise of NEs (Figure S6C), this is also consistent with the experimental data (Figure 3B in (4)).

We also calculate the mean duration time (MDT) of the LTR-off and LTR-on states. Different from the Detailed-Balanced situation, NEs can lengthen the MDT at the LTR-on states more significantly than lengthen the MDT at the LTR-off states (Figure 6F, Figure S7C-F). Further, the effective inhibiting parameter *f*_inh_ ≈ 1 > 0 (See Supplementary Section 2.9 for the exact definition of this effective parameter) means that AC does inhibit NE’s function of reducing the transition rate from the LTR-off states to the LTR-on states (Figure 6F). These simulation results verified the conclusion we made in the LTR-two-state model, that NE can synergize with AC and NE on reactivating latent HIV only when *f*_inh_ > 0. Now we know that AC’s inhibiting NE’s function of reducing effective *k*_on_ is achieved by the clockwise cycle flux driven by the energy input.

However, same as in the LTR-two-state model, Noise Enhancer can amplify Activator’s reactivating latent HIV only if *k*_unbindp_ (equivalent to *k*_off_ in the LTR-two-state model) is greater than 10^−2^ (Figure S8A). To explain this necessary condition, we analyze the timescale of Tat transactivation dynamics and the timescale of LTR transitions. We find that it takes about *τ*_0_ ≈ 20 hours on average for Tat transactivation and then LTR can maintain the activated state for a long time (Figure S8B-C). Therefore, if *k*_unbindp_ is very small compared to the time scale of 1/*τ*_0_, the duration time of LTR-on state without NE presented will be long enough for Tat transactivation with a large possibility, so further reducing *k*_unbindp_ by NE will have little effect (Figure S8B). Only when *k*_unbindp_ is not small compared to 1/*τ*_0_, the duration time of LTR-on state without NE presented is typically not long enough for Tat transactivation. In this case, noise enhancer lengthening the duration time at the LTR-on state will provide Tat more time to reactivate latent HIV (Figure S8B), resulting in the drug synergy with Activator.

Finally, to verify the model applicability, we use the same EITST model to explain other important previous experimental observations (Figure S3A-C) including Tat-transactivation-controlled HIV latency establishment operating autonomously from the host cellular state (31) and the bimodal distribution of phenotype bifurcation (10) (See supplementary Section 2.13 for parameter values). Also, in our EITST model, nonmonotonicity and concavity of the distribution of the dwell time are also observed with large magnitude of energy dissipation (Figure S10F).

## Discussion

Retroviruses are mild at beginning infection. Most replicable retroviruses do not cause cytopathic effects, and infected cells hardly produce a defensive response to such viral replication. Animals rarely experience acute symptoms due to infection with retroviruses, but viremia occurs; the host immune responses reduce the production of the virus. The virus cannot be eradicated even though can be suppressed by the host’s immune response. Therefore, low levels of viremia usually accompany the life of the host (46). As a typical example of the retrovirus, the long-lived latent HIV-1 are the main obstacles to a clinical cure (2). Recently, noise enhanced synergistic combinations of drugs (Activators and Noise Enhancers) were reported more effective than other reactivation cocktails on reactivating HIV latency (4). We here propose an LTR-four-state model with Tat transactivation (the cooperativity is only one) to reveal the mechanism of this synergy, produced by the combination of Activator and Noise Enhancer. By analysis and simulation of the model, we find that the drug synergy on HIV latency reactivation depends the distribution of energy input and the direction of cycle flux of the system. Such a proposed non-equilibrium mechanism should be useful for identifying new drug synergy or regulating the existing drug synergy in HIV and a diverse class of retroviruses infection therapies, most of which have latency states preventing from a complete cure.

Design principle for specific biological functions have been extensively explored, such as reliable cell decisions (47), adaptation (48), robust and tunable biological oscillation (49), and dual function of adaptation and noise attenuation (50). Some of these functions, such as precise biochemical oscillations and accurate adaptation, were found crucially depending on the energy dissipation (26, 27). Here, we show that the drug synergy between NE and AC on reactivating HIV latency also essentially depends on the directional dissipated chemical energy on the HIV LTR-state transition. This non-equilibrium property could also be used as a potential target for lentivirus latency reversal synergetic therapeutic interventions. The optimization principle of energy input distribution for the highest drug synergy might also apply to the network designing.

Our LTR-four-state model is a minimal model in which the effects of AC and NE are distinguished. This LTR-four-state model can be expanded into a more detailed LTR-six-state model where the Tat positive feedback is modeled through Tat binding to LTR and forming two new states, LTR-P-Tat and LTR*-P-Tat as modeled in the previous study (31). In the LTR-six-state model, the same synergy can be predicted (Figure S9, Supplementary Section 2.4). Hence, our uncovered non-equilibrium mechanism of drug synergy is not dependent on the specific Tat positive feedback mechanism.

The synergy that we studied here does not require cooperative binding between the two categories of drugs (AC and NE or NS), which is essentially different from the drug synergy based on classical equilibrium binding mechanism. It might be the reason why the tested drug synergies in *vitro* often fail in *vivo.* The nonequilibrium model proposed here provides a new perspective to understand the drug synergy mechanism.

## Acknowledgments

We thank Profs. Yuhai Tu and Tom Chou for helpful discussions. The work was supported by National Natural Science Foundation of China (11861130351, 11622101, 11622102,11971037).

## Notes

#### Summary of Updates

Supplemental files updated.

