## Supplementary material for "The Nonequilibrium Mechanism of Noise Enhancer synergizing with Activator in HIV Latency Reactivation": SI texts and SI Figures S1-S10

**This PDF file includes:**

Supplementary text
Figures S1 to S9
Tables: Table S1-1 Table S1-2 Table S2-1 Table S2-2 Table S2-3 Table S2-4
SI References

### Contents

1. LTR-2-State Model and Simulation 1
   1.1 LTR-2-state model 1
   1.2 The functions of AC and NE in the LTR-2-state model 1
   1.3 finh, the degree of AC's inhibition upon the reduction of kon induced by NE 2
   1.4 Simulation of reactivation ratio 2
   1.5 The synergy between AC and NE 2
   1.6 Parameter values 3
2. LTR-4-State Model and Simulation 4
   2.1 Detailed Balance LTR-4-state model 4
   2.2 Non-Detailed-Balance LTR-4-state model 5
   2.3 GFP expression without feedback, calculation of mean and noise of LTR 6
   2.4 Tat expression with positive feedback 8
   2.5 Probability of LTR-on states Pon 10
   2.6 Probability flux and cycle flux 10
   2.7 Reactivation ratio 10
   2.8 Mean duration time 10
   2.9 finh, the degree of AC's inhibition upon the reduction of λon induced by NE 11
   2.10 Theorem on the relation between drug synergy of Ponand cyclic probability flux 11
   2.12 Distribution of duration time at LTR-on/off states in equilibrium system 14
   2.13 Parameter values 14
3. Supplemental Figures 16

References 28

### 1. LTR-2-State Model and Simulation

#### 1.1 LTR-2-state model

We employed a well-established LTR-2-state model with Tat positive feedback from previous study (Figure 2A) (1, 2):

$$
\begin{gathered}
LTR_{\text{on}} \xrightarrow{k_{off}} LTR_{\text{off}} \\
LTR_{\text{off}} \xrightarrow{k_{on}} LTR_{\text{on}} \\
LTR_{\text{on}} + Tat \xrightarrow{k_{bind}} LTR_{\text{on-Tat}} \\
LTR_{\text{on-Tat}} \xrightarrow{k_{unbind}} LTR_{\text{on}} + Tat \\
LTR_{\text{on}} \xrightarrow{k_m} mRNA + LTR_{\text{on}} \\
LTR_{\text{on-Tat}} \xrightarrow{k_{transact}} mRNA + LTR_{\text{on-Tat}} \\
mRNA \xrightarrow{k_p} Tat + mRNA \\
mRNA \xrightarrow{d_m} \emptyset \\
Tat \xrightarrow{d_p} \emptyset
\end{gathered}
\tag{S1-1}
$$

In this model, the promoter LTR can toggle between active and inactive states with transition rates $k_{off}$ and $k_{on}$. Tat can bind/unbind to LTR (TAR) with rate $k_{bind}$ and $k_{unbind}$, then transactivate LTR-on state once bound to TAR with a higher transcription rate $k_{transact}$ than the transcription rate $k_m$ at LTR-on state due to Tat's enhancing LTR transcriptional elongation. mRNA can translate into protein at rate $k_p$. Also, mRNA and Tat will degrade at rate $d_m$ and $d_p$ respectively.

#### 1.2 The functions of AC and NE in the LTR-2-state model

For clarity, we defined the rate variables as following:

$k_{on}(k_{off})$: the LTR turning on(off) rate in untreated HIV/LTR-GFP infected cells.

$k_{on,AC}(k_{off,AC})$: the LTR turning on(off) rate with only Activator added to HIV/LTR-GFP infected cells.

$k_{on,NE}(k_{off,NE})$: the LTR turning on(off) rate with only Noise Enhancer added to HIV/LTR-GFP infected cells.

$k_{on,AC,NE}(k_{off,AC,NE})$: the LTR turning on(off) rate with both Activator and Noise Enhancer added to HIV/LTR-GFP infected cells.

$k_{on,NS}$, $k_{off,NS}$, $k_{on,AC,NS}$ and $k_{off,AC,NS}$ are defined in the same way.

In this LTR-2-state model, the functions of AC and NE are assumed as following (1): adding AC increases $k_{on}$ to $k_{on}r_{AC}$, while adding NE reduces $k_{on}$ and $k_{off}$ to $k_{on}r_{NE}$ and $k_{off}r_{NE}$ ($r_{NE} < 1$), respectively, with their ratio fixed. From these assumptions, we have:

$$
\begin{cases} k_{on,AC} = k_{on}r_{AC} \\ k_{off,AC} = k_{off} \end{cases}
\tag{S1-2}
$$

and

$$\begin{cases} k_{on,NE} = k_{on} r_{NE} \\ k_{off,NE} = k_{off} r_{NE} \end{cases} \tag{S1-3}$$

The values of $k_{on,AC,NE}$ and $k_{off,AC,NE}$ will be discussed in the next section.

#### 1.3 $f_{inh}$, the degree of AC's inhibition upon the reduction of $k_{on}$ induced by NE

If the functions of AC and NE are working separately and independently, then

$$\begin{cases} k_{on,AC,NE} = k_{on} r_{AC} r_{NE} \\ k_{off,AC,NE} = k_{off} r_{NE} \end{cases}$$

However, there might be some interaction between AC and NE's function. For AC and NE to has synergy on HIV latency reactivation, AC might inhibit NE's function on $k_{on}$. We quantified the inhibition of AC on NE's function on $k_{on}$ as:

$$f_{inh} = 1 - \log_{r_{NE}}\left(\frac{k_{on,AC,NE}}{k_{on} r_{AC}}\right) = \frac{\ln(k_{on,AC,NE}) - \ln(k_{on,AC})}{\ln(k_{off,AC}) - \ln(k_{off,AC,NE})} + 1 \tag{S1-4}$$

Then we have:

$$\begin{cases} k_{on,AC,NE} = k_{on} r_{AC} r_{NE}^{1-f_{inh}} \\ k_{off,AC,NE} = k_{off} r_{NE} \end{cases} \tag{S1-5}$$

$$f_{inh} \in [0,1]$$

$f_{inh} = 0$ represents AC does not inhibit NE's function of reducing $k_{on}$ (Figure S2C, left panel), and $f_{inh} > 0$ means that AC does inhibit NE's function of reducing $k_{on}$. Particularly, $f_{inh} = 1$ means that NE's function of reducing $k_{on}$ is completely inhibited by AC (Figure S2C, right panel).

#### 1.4 Simulation of reactivation ratio

The stochastic LTR-2-state model coupled with Tat positive feedback (Eqs (S1-1)) was simulated using the Stochastic Simulation Algorithm(SSA) (3).

Reactivation Ratio is the ratio of trajectory numbers with activated HIV (#Tat > 75) up to 100 hours and the total number of trajectories starting from the latency state ($LTR_{off} = 1$, copy numbers of all other species=0, simulated 5000~10000 cells) at time 0h. These simulations were implemented via Matlab™ with the parameters shown in Table S1-1 and Table S1-2.

#### 1.5 The synergy between AC and NE

From the experiments, the drug synergy is that NE can significantly amplify the reactivation of latent HIV caused by AC, while NE itself cannot reactivate latent HIV, i.e. 1+1>2 (Figure 1B) (1). Mathematically, we define the synergy on reactivating latent HIV as:

$$\text{Synergy} = R_{AC,NE} - R_{AC} \tag{S1-6}$$

$R_{AC,NE}$ is the reactivation ratio of the latent HIV under parameter $k_{on,AC,NE}$ and $k_{off,AC,NE}$, corresponding to adding

AC and NE together; $R_{AC}$ is the reactivation ratio of the latent HIV under parameter $k_{on,AC}$ and $k_{off,AC}$, corresponding to adding only AC. The calculation of reactivation ratio has been explained in section 1.4.

#### 1.6 Parameter values

Table S1-1. Parameter values in Figure 2, Figure S2

| Parameter | Value |
|---|---|
| $k_{on}$ | varied (for Figure S2G, $k_{on} = 0.0001$) |
| $k_{off}$ | Varied (for Figure S2G-I, $k_{off} = 0.01$) |
| ($r_{NE}$) | (0.5 for Figure 2C-D) |
| $k_m$ | 1 |
| $d_m$ | 1 |
| $k_p$ | 10 |
| $k_{bind}$ | 0.01 |
| $k_{unbind}$ | 0.01 |
| $k_{transact}$ | 5 |
| $d_p$ | 0.125 |
| Total Cell Number | 10000 |
| Time steps (Iterations) | 50000 |
| Observation time | 100 hours |
| Threshold for reactivation | 75 |

All the parameter values except $k_{on}$ and $k_{off}$ are the same as those in (2) which is quantified by single-cell analysis (4-6). $k_{on}$ and $k_{off}$ mainly depends on the integration sites of HIV in human genome. Thus, the values of $k_{on}$ and $k_{off}$ can vary in a large range. In addition, we want to investigate how the parameter values of $k_{on}$ and $k_{off}$ infect the synergy. It is necessary to vary the values of parameter $k_{on}$ and $k_{off}$. Based on the previous study (2), we set the range of $k_{on}$ and $k_{off}$ from $10^{-4}$ to $10^{0}$.

Table S1-2. Parameter values in Figure 2E-G

| Parameter | Figure 2E | Figure 2F | Figure 2G |
|---|---|---|---|
| $k_{on,AC}$ | 0.0008 | 0.008 | 0.08 |
| $k_{off,AC}$ | 0.008 | 0.08 | 0.8 |
| $k_{off,AC,NS}$ | 0.002 | 0.02 | 0.2 |
| $k_{off,AC,NE}$ | 0.032 | 0.32 | 3.2 |

### 2. LTR-4-State Model and Simulation

#### 2.1 Detailed Balance LTR-4-state model

We built a LTR-4-state model under detailed balance (Figure 3A):

$$\begin{gathered}
LTR \xrightarrow{k_{act}\gamma} LTR^* \\
LTR^* \xrightarrow{k_{unact}} LTR \\
LTR^* \xrightarrow{\omega k_{bindp}} LTR^*P \\
LTR^*P \xrightarrow{k_{unbindp}} LTR^* \\
LTR^*P \xrightarrow{k_{unact}} LTRP \\
LTRP \xrightarrow{\omega k_{act}\gamma} LTR^*P \\
LTRP \xrightarrow{k_{unbindp}e^{-\alpha}} LTR \\
LTR \xrightarrow{k_{bindp}e^{-\alpha}} LTRP
\end{gathered} \tag{S2-1}$$

In our model, there are four different promotor states: LTR is the free state, LTR* is the activated state but without RNAP binding; LTR-P and LTR*-P are the corresponding RNAP-bounded states. AC is assumed to promote LTR transiting to the activated state LTR*, e.g. LTR bound with NF-κB, and LTR* recruits RNA polymerase much easier than LTR itself, e.g. the NF-κB bound to LTR acting as a Transcription Factor to recruit RNA polymerase to LTR(7). It has been shown that the screened Noise Enhancer has no effect on post transcription (1), and some NE can increase transcription factors in cells, such as SP1 (8, 9). Similar to (1), we assume that NE, once present, can slows down the switching rates between LTR and LTR-P.

We used Markov jumping process to model the transition among LTR states with AC and/or NE added (Figure 3A). The four states can mutually transit. We assume $S = \{LTR, LTR^*, LTRP, LTR^*P\}$, use $R$ represents $LTR$ state, $R^*$ represents $LTR^*$ state, $R^*P$ represents $LTR^*P$ state, $P$ represents $LTRP$ state. Then $S = \{R, R^*, P, R^*P\}$. We denote that $ON = \{P, R^*P\}$, $OFF = \{R, R^*\}$. The generator matrix (transition rate matrix) is:

$$Q = \begin{bmatrix} k_R & k_{R,R^*} & k_{R,RP} & 0 \\ k_{R^*,R} & k_{R^*} & 0 & k_{R^*,R^*P} \\ k_{P,R} & 0 & k_P & k_{P,R^*P} \\ 0 & k_{R^*P,R^*} & k_{R^*P,P} & k_{R^*P} \end{bmatrix} \tag{S2-2}$$

$$k_i = -\sum_j k_{i,j} \quad i,j \in S$$

Then we can calculate invariant distribution $\pi$:

$$\begin{bmatrix} \pi_R \\ \pi_{R^*} \\ \pi_P \\ \pi_{R^*P} \end{bmatrix}^T \begin{bmatrix} k_R & k_{R,R^*} & k_{R,P} & 0 \\ k_{R^*,R} & k_{R^*} & 0 & k_{R^*,R^*P} \\ k_{P,R} & 0 & k_P & k_{P,R^*P} \\ 0 & k_{R^*P,R^*} & k_{R^*P,P} & k_{R^*P} \end{bmatrix} = \begin{bmatrix} 0 \\ 0 \\ 0 \\ 0 \end{bmatrix}^T \tag{S2-3}$$

More specifically, here we assume in the absence of AC, RNAP binds to LTR at a relatively slow rate $k_{bindp}[P]$ and unbinds fast at rate $k_{unbindp}$; LTR transit to LTR* state with an extremely slow rate $k_{act}$ without AC, but at a much higher rate $k_{act}\gamma$ ($\gamma \gg 1$) once AC is present; RNAP is attracted to bind to LTR* at a higher rate $\omega k_{bindp}$, where $\omega$ is the cooperative interaction factor ($\omega > 1$); when NE is added, LTR will bind and unbind RNAP at a slower rate ($k_{bindp}e^{-\alpha}$, $k_{unbindp}e^{-\alpha}$) with a reduction parameter $\alpha > 0$; LTR-P and LTR*-P can mutually transit at rate $\omega k_{act}\gamma$ and rate $k_{unact}$.

Here, there is no external energy input; it is under detailed balance condition:

$$k_{R,R^*}k_{R^*,R^*P}k_{R^*P,P}k_{P,R} = k_{R,P}k_{P,R^*P}k_{R^*P,R^*}k_{R^*,R} \quad \text{(S2-4)}$$

where $k_{R,R^*} = k_{act}\gamma$, $k_{R^*,R^*P} = \omega k_{bindp}$, $k_{R^*P,P} = k_{unact}$, $k_{P,R} = k_{unbindp}e^{-\alpha}$, $k_{R,P} = k_{bindp}e^{-\alpha}$, $k_{P,R^*P} = \omega k_{act}\gamma$, $k_{R^*P,R^*} = k_{unbindp}$, $k_{R^*,R} = k_{unact}$.

#### 2.2 Non-Detailed-Balance LTR-4-state model

Breaking the detailed balance condition in the Detailed Balance LTR-4-state model, we can built non-Detailed-Balance LTR-4-state models. We first built it with energy input only through a single transition: Energy input can influence any single transition rate in the Detailed Balance LTR-4-state model through multiplying it by a factor of $e^{\beta}$. Such an energy input will cause clockwise (c.w.) probability flux or counter-clockwise (c.c.w.) probability flux. The details of the non-Detailed-Balance model are listed in Table S2-1.

We also built an EITST(Energy Input on the Two Specific Transition rates) LTR-4-state model, in which part of the energy input is through reducing the transition rate from LTR*-P to LTR* by multiplying $e^{-\beta_2}$, and the other half is through increasing the transition rate from LTR*-P to LTR-P by multiplying $e^{\beta_1}$ (Figure 6A), instead of with energy input on a single transition. The details of the EITST model are shown below.

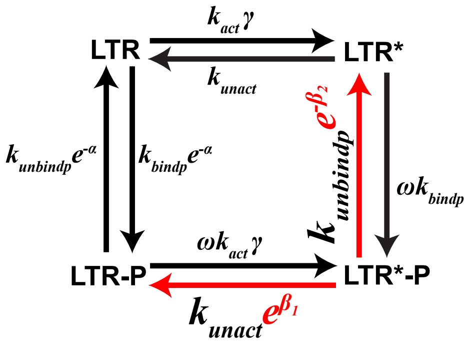

Figure. The rate formula in EITST model

Table S2-1. The red arrow indicates the single transition rate changed by the energy input $e^{\beta}$.

c.w.=clockwise, c.c.w.=counter-clockwise

| Model label | Schematic of model | Sign of energy | Direction of probability flux |
| --- | --- | --- | --- |
| Detailed Balance | 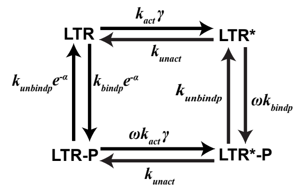 | $\beta = 0$ | No Flux |
| R → R* | 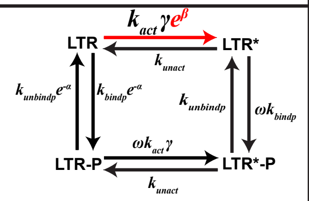 | $\beta > 0$ | c.w. |
| | | $\beta < 0$ | c.c.w. |
| R* → R*P | 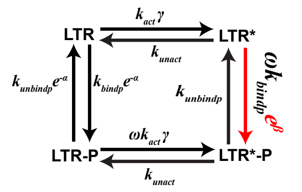 | $\beta > 0$ | c.w. |
| | | $\beta < 0$ | c.c.w. |
| R*P → P | 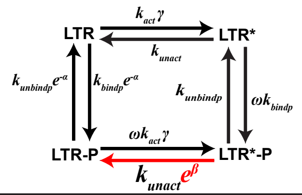 | $\beta > 0$ | c.w. |
| | | $\beta < 0$ | c.c.w. |
| P → R | 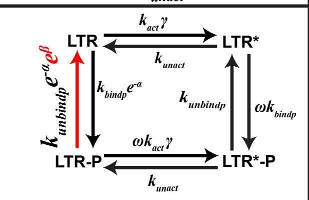 | $\beta > 0$ | c.w. |
| | | $\beta < 0$ | c.c.w. |

| Model label | Schematic of model | Sign of energy | Direction of probability flux |
| --- | --- | --- | --- |
| R* → R | 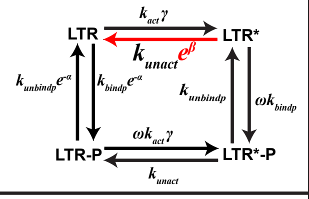 | $\beta > 0$ | c.c.w. |
| | | $\beta < 0$ | c.w. |
| R*P → R* | 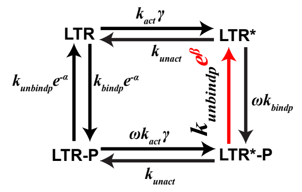 | $\beta > 0$ | c.c.w. |
| | | $\beta < 0$ | c.w. |
| P → R*P | 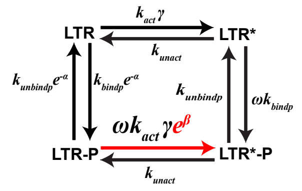 | $\beta > 0$ | c.c.w. |
| | | $\beta < 0$ | c.w. |
| R → P | 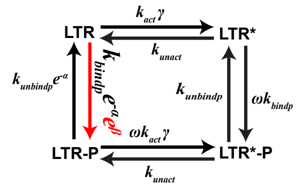 | $\beta > 0$ | c.c.w. |
| | | $\beta < 0$ | c.w. |

### 2.3 GFP expression without feedback, calculation of mean and noise of LTR

We describe the dynamics of LTR-GFP vector expression using the following chemical reactions (combined with reactions (S2-1)):

$$LTR^*P \xrightarrow{k_m} mRNA + LTR^*P$$
$$LTRP \xrightarrow{k_m} mRNA + LTRP$$
$$mRNA \xrightarrow{k_p} GFP + mRNA \qquad \text{(S2-5)}$$
$$mRNA \xrightarrow{d_m} \emptyset$$
$$GFP \xrightarrow{d_p} \emptyset$$

For the LTR-GFP vector, with RNAP bond to LTR (i.e. LTR-P state or LTR*-P state), the downstream DNA of

LTR can be transcribed into mRNA at rate $k_m$ and then translate into protein at rate $k_{GFP}$. Also, mRNA and GFP will degrade at rate $d_m$ and $d_{GFP}$ respectively (Figure 3B). To Calculate the Noise and Mean of GFP of this system, we need calculate the first and second moment of GFP:

$$< GFP > (t) = \sum_{i \in S} \sum_{m=0}^{\infty} \sum_{n=0}^{\infty} nP(LTR = i, mRNA = m, GFP = n, t)$$

and

$$< GFP^2 > (t) = \sum_{i \in S} \sum_{m=0}^{\infty} \sum_{n=0}^{\infty} n^2 P(LTR = i, mRNA = m, GFP = n, t)$$

Here, $P(LTR = i, mRNA = m, t)$ represents the probability of LTR staying at $i$ state and $\#mRNA = m$ at time $t$; $P(LTR = i, mRNA = m, GFP = n)$ represents the probability of LTR staying at $i$ state and $\#mRNA = m$ and $\#GFP = n$ at time $t$. For the following, when the variable/quantity/moment is not written as an explicit function of time $t$, it mean the steady state value. Then we sum up the related master equation and calculate the steady state (the derivative is zero):

$$< GFP >= \frac{k_p}{d_p} < mRNA > \tag{S2-6}$$

and

$$< GFP^2 >= \frac{k_p}{d_p} (< GFP\ mRNA > + < mRNA >) \tag{S2-7}$$

To calculate the above quantity, we need calculate the following moments of mRNA and GFP:

$$< mRNA > (t) = \sum_{i \in S} \sum_{m=0}^{\infty} mP(LTR = i, mRNA = m, t)$$

$$< mRNA^2 > (t) = \sum_{i \in S} \sum_{m=0}^{\infty} m^2 P(LTR = i, mRNA = m, t)$$

$$< GFP\ mRNA > (t) = \sum_{i \in S} \sum_{m=0}^{\infty} \sum_{n=0}^{\infty} mnP(LTR = i, mRNA = m, GFP = n, t)$$

and similarly, we sum up the related master equation and calculate the steady state (the derivative is zero), followed by

$$< mRNA >= \frac{1}{d_m} \sum_{i \in S} k_{m,i} \pi_i \tag{S2-8}$$

$$< GFP\ mRNA >= \frac{\sum_{i \in S} k_{m,i} < GFP >_i + k_p < mRNA^2 >}{d_m + d_p} \tag{S2-9}$$

$$< mRNA^2 >= \frac{1}{d_m} \sum_{i \in S} k_{m,i} < mRNA >_i + \frac{1}{d_m} \sum_{i \in S} k_{m,i} \pi_i \tag{S2-10}$$

where

$$k_{m,i} = \begin{cases} k_m, i \in ON \\ 0, i \in OFF \end{cases}$$

$$< mRNA >_i = \sum_{m=0}^{\infty} mP(LTR = i, mRNA = m) \tag{S2-11}$$

$$< GFP >_i = \sum_{n=0}^{\infty} n \sum_{m=0}^{\infty} P(LTR = i, mRNA = m, GFP = n)$$

for all $i \in S$. $< mRNA >_i$ and $< GFP >_i$ satisfy the linear equations (the steady state of Master Equations):

$$(d_m I_4 - Q)\begin{bmatrix} < mRNA >_R \\ < mRNA >_{R^*} \\ < mRNA >_P \\ < mRNA >_{R^*P} \end{bmatrix} = \begin{bmatrix} k_{m,R}\pi_R \\ k_{m,R^*}\pi_{R^*} \\ k_{m,P}\pi_P \\ k_{m,R^*P}\pi_{R^*P} \end{bmatrix} \quad \text{(S2-12)}$$

$$(d_p I_4 - Q)\begin{bmatrix} < GFP >_R \\ < GFP >_{R^*} \\ < GFP >_P \\ < GFP >_{R^*P} \end{bmatrix} = k_p \begin{bmatrix} < mRNA >_R \\ < mRNA >_{R^*} \\ < mRNA >_P \\ < mRNA >_{R^*P} \end{bmatrix} \quad \text{(S2-13)}$$

where $I_4$ is the $4 \times 4$ identity matrix, $Q$ is the generator matrix for the LTR-4-state model.

We solved the linear equations (S2-12) (S2-13), then substituted $< mRNA >_i$ and $< GFP >_i$ for $i \in S$ into equations (S2-9) (S2-10). We then substituted (S2-8) (S2-9) (S2-10) into equations (S2-6) (S2-7). Using the above calculation and submission, we have the Noise of LTR-GFP vector, $\frac{<GFP^2>-<GFP>^2}{<GFP>^2}$, and the mean of LTR-GFP vector, $< GFP >$.

### 2.4 Tat expression with positive feedback

We describe the dynamics of the full length HIV vector expression with Tat positive feedback using the following chemical reactions (combined with reactions (S2-1)):

$$\begin{gathered} LTR^*P \xrightarrow{k_m} mRNA + LTR^*P \\ LTRP \xrightarrow{k_m} mRNA + LTRP \\ mRNA \xrightarrow{k_{Tat}} Tat + mRNA \\ mRNA \xrightarrow{d_m} \emptyset \\ Tat \xrightarrow{d_{Tat}} \emptyset \end{gathered} \quad \text{(S2-14)}$$

The Tat forms positive feedback by enhancing the elongation of initial transcribed mRNA of HIV (10, 11) and by stabilizing the HIV activation (2). We model these two function by following:

$$k_m = k_{mbasal} + k_{trs1}\frac{\frac{Tat}{k_{trs2}}}{1+\frac{Tat}{k_{trs2}}} \quad \text{(S2-15)}$$

and

$$k_{unbindp} = \frac{K_{threshold}^3 + \delta Tat^3}{K_{threshold}^3 + Tat^3} k_{unbindp0} \quad \text{(S2-16)}$$

All parameter values are shown in Table S2-3.

The corresponding ordinary differential equations of Tat protein and mRNA are at LTR-on states are:

$$\begin{cases} \frac{dmRNA}{dt} = k_{mbasal} + k_{trs1}\frac{\frac{Tat}{k_{trs2}}}{1+\frac{Tat}{k_{trs2}}} - d_m mRNA \\ \frac{dTat}{dt} = k_{Tat}mRNA - d_{Tat}Tat \end{cases}$$

To prove the results is independent from the specific details of the model, we also applied another model adapted from (2) shown below.

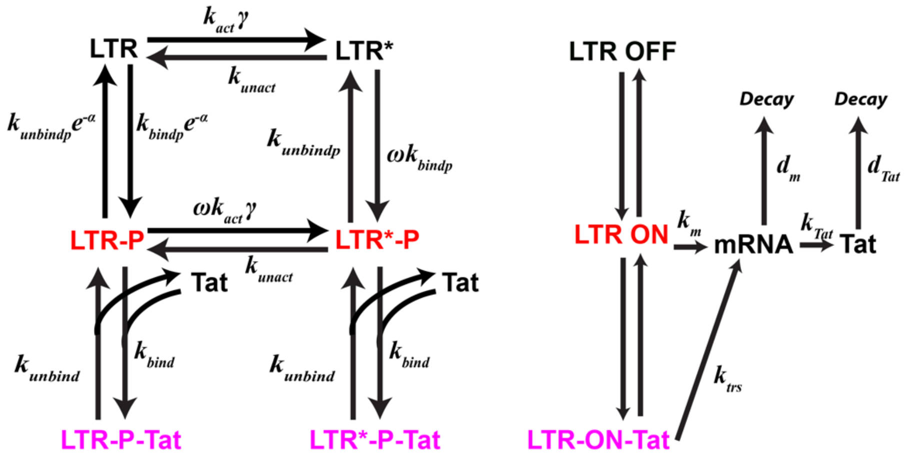

In this model, similar to LTR-2-state Tat positive feedback model, the Tat forms positive feedback through binding to LTR(TAR) Tat can bind/unbind to LTR(TAR) with rate $k_{bind}$ and $k_{unbind}$, then transactivate LTR-on state once bound to TAR with a much higher transcription rate $k_{trs}$ than the transcription rate $k_m$ at LTR-on state due to Tat's enhancing LTR transcriptional elongation. The system can be expressed by the following chemical reactions (combined with reactions (S2-1)):

$$\begin{gathered} LTRP + Tat \xrightarrow{k_{bind}} LTRP_{Tat} \\ LTRP_{Tat} \xrightarrow{k_{unbind}} LTRP + Tat \\ LTR^*P + Tat \xrightarrow{k_{bind}} LTR^*P_{Tat} \\ LTR^*P_{Tat} \xrightarrow{k_{unbind}} LTR^*P + Tat \\ LTRP \xrightarrow{k_m} mRNA + LTRP \\ LTR^*P \xrightarrow{k_m} mRNA + LTR^*P \\ LTRP_{Tat} \xrightarrow{k_{trs}} mRNA + LTRP_{Tat} \\ LTR^*P_{Tat} \xrightarrow{k_{trs}} mRNA + LTR^*P_{Tat} \\ mRNA \xrightarrow{k_{Tat}} Tat + mRNA \\ mRNA \xrightarrow{d_m} \emptyset \\ Tat \xrightarrow{d_{Tat}} \emptyset \end{gathered} \qquad \text{(S2-17)}$$

All parameter values are the same as LTR-2-state Tat positive feedback model, shown in Table S1-1.

### 2.5 Probability of LTR-on states $P_{on}$

From (S2-3), we have invariant distribution of LTR-4-state model, $\pi_i$ for $i \in S$. We then calculated the probability of LTR-on states (LTR-P state and LTR*-P state):

$$P_{on} = \pi_P + \pi_{R^*P} \quad \text{(S2-18)}$$

### 2.6 Probability flux and cycle flux

From invariant distribution (S2-3) and the transition rates, we can calculate the probability flux of LTR-4-state model at steady state:

$$J_{ij} = \pi_i k_{i,j}$$

for $i, j \in S$. And net flux from state I to state j is defined as $J_{ij} - J_{ji}$.

In such a 4-state model, there is only one cycle (LTR->LTR*->LTR*-P->LTR-P->LTR), which is clockwise, and its reversed one. The net cycle flux of the clockwise cycle is $J_c = J_{R,R*} = J_{R*,R*P} = J_{R*P,RP} = J_{RP,R}$, and the net flux of the reversed counterclockwise cycle is $-J_c$.

### 2.7 Reactivation ratio

The stochastic LTR-4-state model coupled with Tat positive feedback (Eqs (S2-1) (S2-14) (S2-15) (S2-16)) was simulated using the Stochastic Simulation Algorithm(SSA), or say 'Gillespie' algorithm (3), because of the difficulty to analytically calculate the model with feedback.

Reactivation Ratio is the ratio of activated HIV (#Tat > 75) trajectory number up to 100 hour and the total number of trajectories starting from latent state (LTR=1, copy numbers of all other species=0, simulated 5000~10000 cells) at time 0h. These simulations were implemented via Matlab™ with the parameters shown in Table S2-2 and Table S2-3.

### 2.8 Mean duration time

We need a theorem to calculate mean duration time.

**Theorem** (12)

Let $\{X_t, t \geq 0\}$ be a continuous time Markov chain on state space $S$, with generator matrix $Q = (q_{ij})$. $S_1$ and $S_2$ are subspaces of $S$ satisfying:

$$S = S_1 \cup S_2,\ S_1 \cap S_2 = \emptyset,\ S_1 \neq \emptyset,\ S_2 \neq \emptyset.$$

Suppose the invariant distribution $\mu = \{\mu_i : i \in S\}$ exists, then the mean duration in $S_1$ (denoted by $\tau$) takes the form of

$$\tau = \frac{\sum_{i \in S_1} \mu_i}{\sum_{i \in S_1} \sum_{j \in S_2} \mu_i q_{ij}}.$$

See (12) for proof.

For the LTR-4-state continuous time Markov chain with transitions showed in Eqs (S2-1), the generator $Q$ is

(S2-2), and the invariant distribution $\pi_i$ for $i \in S$ can be derived from linear Eqs (S2-3). We assume that $S_1 = ON = \{LTRP, LTR^*P\}$, $S_2 = OFF = \{LTR, LTR^*\}$. We define $\tau_{ON}(\tau_{OFF})$ as the mean duration time of LTR stay at ON(OFF) states. By the theorem, we have:

$$\tau_{ON} = \frac{\sum_{i \in ON} \pi_i}{\sum_{i \in ON} \sum_{j \in OFF} \pi_i q_{ij}} \quad \text{(S2-19)}$$

$$\tau_{OFF} = \frac{\sum_{i \in OFF} \pi_i}{\sum_{i \in OFF} \sum_{j \in ON} \pi_i q_{ij}} \quad \text{(S2-20)}$$

### 2.9 $f_{inh}$, the degree of AC's inhibition upon the reduction of $\lambda_{on}$ induced by NE

We regarded the reciprocal of the Mean Duration Time as the transition rates between LTR-on and LTR-off states, $\lambda_{on}$ and $\lambda_{off}$,

$$\begin{cases} \lambda_{on} = \frac{1}{\tau_{OFF}} \\ \lambda_{off} = \frac{1}{\tau_{ON}} \end{cases} \quad \text{(S2-21)}$$

which are equivalent to $k_{on}$ and $k_{off}$ in the effective LTR-2-State model. Then we defined the effective $f_{inh}$ using the formula (S1-4), i.e.

$$f_{inh} = \frac{\ln(\lambda_{on,AC,NE}) - \ln(\lambda_{on,AC})}{\ln(\lambda_{off,AC}) - \ln(\lambda_{off,AC,NE})} + 1 \quad \text{(S2-22)}$$

$\lambda_{on,AC}$ and $\lambda_{off,AC}$ correspond to the LTR-state model with only AC added, i.e. $\gamma \gg 1$, $\alpha = 0$; $\lambda_{on,AC,NE}$ and $\lambda_{off,AC,NE}$ correspond to the LTR-state model with both AC and NE added, i.e. $\gamma \gg 1$, $\alpha > 0$. Similar we can define $\lambda_{on,AC,NS}$ and $\lambda_{off,AC,NS}$.

### 2.10 Theorem on the relation between drug synergy of $P_{on}$ and cyclic probability flux

Generally, our non-detailed-balanced LTR-4-state model can be described as below:

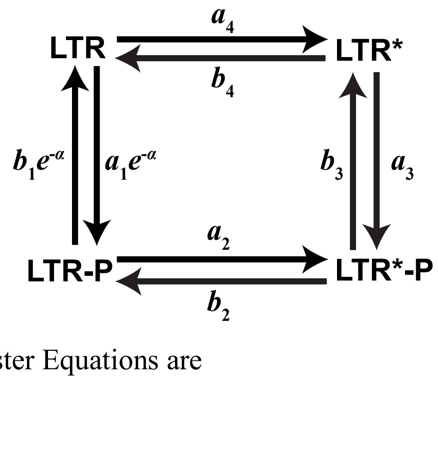

The corresponding Chemical Master Equations are

$$\begin{cases} \frac{dP_{LTR}}{dt} = -(a_1e^{-\alpha} + a_4)P_{LTR} + b_1e^{-\alpha}P_{LTRP} + b_4P_{LTR^*} \\ \frac{dP_{LTRP}}{dt} = a_1e^{-\alpha}P_{LTR} - (b_1e^{-\alpha} + a_2)P_{LTRP} + b_2P_{LTR^*P} \\ \frac{dP_{LTR^*P}}{dt} = a_2P_{LTRP} - (b_2 + b_3)P_{LTR^*P} + a_3P_{LTR^*} \\ \frac{dP_{LTR^*}}{dt} = a_4P_{LTR} + b_3P_{LTR^*P} - (b_4 + a_3)P_{LTR^*} \end{cases} \tag{S2-23}$$

At steady state, the derivative is zero. Then, with the net flux $J$ introduced, the equations above at steady state can be simplified to a set of equations:

$$\begin{cases} J = -a_1e^{-\alpha}P_{LTR} + b_1e^{-\alpha}P_{LTRP} \\ J = -a_2P_{LTRP} + b_2P_{LTR^*P} \\ J = -b_3P_{LTR^*P} + a_3P_{LTR^*} \\ J = -b_4P_{LTR^*} + a_4P_{LTR} \end{cases} \tag{S2-24}$$

Here, $J > 0$ indicates a clockwise cycle net flux while $J < 0$ means a counter-clockwise one. And note that **synergy** of $P_{on}$ is defined as the increase of $P_{on}(\alpha) = P_{LTRP}(\alpha) + P_{LTR^*P}(\alpha)$ when there exists NE ($\alpha > 0$) and the decrease of $P_{on}(\alpha)$ when NS is present ($\alpha < 0$), compared with $P_{on}$ in the merely-AC case ($\alpha = 0$)

**Theorem 1.** For any positive $a_i$ and $b_i$ $(i = 1, 2, 3, 4)$,

(1) If $J > 0$, then $P_{on}(\alpha) > P_{on}(0)$ for $\alpha > 0$ and $P_{on}(\alpha) < P_{on}(0)$ for $\alpha < 0$;

(2) If $J < 0$, then $P_{on}(\alpha) < P_{on}(0)$ for $\alpha > 0$ and $P_{on}(\alpha) > P_{on}(0)$ for $\alpha < 0$.

**Proof.** We begin with the case of $\alpha = 0$. From the kinetics equations (S2-24), we can eliminate the probabilities at the OFF state:

$$P_{LTR} = -\frac{J}{a_1} + \frac{b_1}{a_1}P_{LTRP},$$
$$P_{LTR^*} = -\frac{a_1 + a_4}{a_1b_4}J + \frac{a_4b_1}{a_1b_4}P_{LTRP}.$$

Therefore,

$$\left(1 + \frac{a_3(a_1 + a_4)}{a_1b_4}\right)J = -b_3P_{LTR^*P} + \frac{a_3a_4b_1}{a_1b_4}P_{LTRP},$$
$$J = b_2P_{LTR^*P} - a_2P_{LTRP},$$

and

$$P_{LTRP} = \frac{a_1b_4(b_2 + b_3) + a_3b_2(a_1 + a_4)}{a_3a_4b_1b_2 - a_1a_2b_3b_4}J,$$
$$P_{LTR^*P} = \frac{a_1a_2(a_3 + b_4) + a_3a_4(a_2 + b_1)}{a_3a_4b_1b_2 - a_1a_2b_3b_4}J.$$

Since $P_{LTR} + P_{LTR^*} + P_{LTRP} + P_{LTR^*P} = 1$ gives that

$$-\frac{J}{a_1} + \frac{b_1}{a_1}P_{LTRP} - \frac{a_1 + a_4}{a_1b_4}J + \frac{a_4b_1}{a_1b_4}P_{LTRP} + P_{LTRP} + P_{LTR^*P} = 1,$$

we have

$$-\frac{a_1 + a_4 + b_4}{a_1b_4} + \frac{a_4b_1 + b_1b_4 + a_1b_4}{a_1b_4}\frac{a_1b_4(b_2 + b_3) + a_3b_2(a_1 + a_4)}{a_3a_4b_1b_2 - a_1a_2b_3b_4}$$

$$+\frac{a_1a_2(a_3+b_4)+a_3a_4(a_2+b_1)}{a_3a_4b_1b_2-a_1a_2b_3b_4}=\frac{1}{J}.$$

By replacing $\mathrm{a}_1$ and $b_1$ with $\mathrm{a}_1e^{-\alpha}$ and $b_1e^{-\alpha}$, we obtain the general equality:

$$a_1(a_2a_3+a_2b_3+a_2b_4+a_3b_2+b_2b_4+b_3b_4)+e^{\alpha}(a_2a_3a_4+a_2a_4b_3+a_2b_3b_4+a_3a_4b_2)$$
$$+b_1(a_3a_4+a_3b_2+a_4b_2+a_4b_3+b_2b_4+b_3b_4)=\frac{a_3a_4b_1b_2-a_1a_2b_3b_4}{J(\alpha)}.$$

It is easy to see that:

(1) $J(\alpha)$ decreases with $\alpha$ when $J>0$ and increases with $\alpha$ when $J<0$, i.e.

$$J(0)-J(\alpha)>0,\ \text{if } J>0 \tag{S2-25}$$

$$J(0)-J(\alpha)<0,\ \text{if } J<0 \tag{S2-26}$$

(2) $e^{\alpha}J(\alpha)$ increases with $\alpha$ when $J>0$ and decreases with $\alpha$ when $J<0$;

$$e^{\alpha}J(\alpha)-J(0)>0,\ \text{if } J>0 \tag{S2-27}$$

$$e^{\alpha}J(\alpha)-J(0)<0,\ \text{if } J<0 \tag{S2-28}$$

(3) $J(\alpha)$ shares the same sign with $J(0)$.

$$J(\alpha)J(0)>0 \tag{S2-29}$$

When NE/NS concentration approaches zero which means $\alpha=0$, our model reduces to the general equality in (Jia, 2012):

$$P_{\text{on}}(\alpha)-P_{\text{on}}^{0}=(P_{\text{on}}^{\infty}-P_{\text{on}}^{0})P_{\text{on}}(\alpha)-\left(\frac{P_{\text{on}}^{\infty}}{a_3}-\frac{P_{\text{on}}^{0}}{a_1}\right)J, \tag{S2-30}$$

where $P_{\text{on}}^{0}=\frac{a_1}{a_1+b_1}$, $P_{\text{on}}^{\infty}=\frac{a_3}{a_3+b_3}$ remain the same under any value of α. From (S2-30), we have:

$$(1-P_{\text{on}}^{\infty}+P_{\text{on}}^{0})P_{\text{on}}(\alpha)=P_{\text{on}}^{0}-\left(\frac{P_{\text{on}}^{\infty}}{a_3}-\frac{P_{\text{on}}^{0}}{a_1}\right)J, \tag{S2-31}$$

To calculate the synergy $P_{on}(\alpha)-P_{on}(0)$, we subtract equation (S2-31) with $\alpha=0$ from equation (S2-31) with $\alpha>0$ (NE), and we have:

$$(1-P_{\text{on}}^{\infty}+P_{\text{on}}^{0})\big(P_{\text{on}}(\alpha)-P_{\text{on}}(0)\big)=-\left(\frac{1}{a_3+b_3}-\frac{e^{\alpha}}{a_1+b_1}\right)J(\alpha)+\left(\frac{1}{a_3+b_3}-\frac{1}{a_1+b_1}\right)J(0)$$
$$=\frac{J(0)-J(\alpha)}{a_3+b_3}+\frac{e^{\alpha}J(\alpha)-J(0)}{a_1+b_1}. \tag{S2-32}$$

Since $1-P_{\text{on}}^{\infty}+P_{\text{on}}^{0}>0$, and for clockwise cyclic probability flux ($\mathrm{J}>0$), substitute (S2-25) (S2-27) into (S2-32), we can prove that such case predicts positive synergy.

Similarly, for clockwise cyclic probability flux ($\mathrm{J}>0$), for the effect of NS ($\alpha<0$),

$$(1-P_{\text{on}}^{\infty}+P_{\text{on}}^{0})\big(P_{\text{on}}(\alpha)-P_{\text{on}}(0)\big)=\frac{J(0)-J(\alpha)}{a_3+b_3}+\frac{e^{\alpha}J(\alpha)-J(0)}{a_1+b_1}<0$$

which shows that the LTR-4-state model with clockwise cyclic probability flux distinguishes NE and NS well.

Since the monotonicity of both $J(\alpha)$ and $e^{\alpha}J(\alpha)$ gets reverse, by the same token we can prove that for counter-clockwise cyclic probability flux ($J<0$), $P_{\text{on}}(\alpha)-P_{\text{on}}(0)<0$ and no synergy is predicted when $\alpha>0$ (NE), while $P_{\text{on}}(\alpha)-P_{\text{on}}(0)>0$ when $\alpha>0$ (NS).

### 2.12 Distribution of duration time at LTR-on/off states in equilibrium system

According to Tu (13), the distribution of duration time at LTR-on/off states in equilibrium system should be monotonically decreasing and convex, while in the non-equilibrium system these features can be violated. We use the following equations to calculate the distribution of duration time at LTR-off states:

$$\frac{dQ(LTR,\tau)}{d\tau} = k_{R^*,R}Q(LTR^*,\tau) - k_{R,R^*}Q(LTR,\tau) - k_{R,P}Q(LTR,\tau)$$

$$\frac{dQ(LTR^*,\tau)}{d\tau} = -k_{R^*,R}Q(LTR^*,\tau) + k_{R,R^*}Q(LTR,\tau) - k_{R^*,R^*P}Q(LTR^*,\tau)$$

With initial conditions:

$$Q(LTR,0) = A_{off}k_{P,R}\pi_P$$

$$Q(LTR^*,0) = A_{off}k_{R^*P,R^*}\pi_{R^*P}$$

Where

$$A_{off} = \frac{1}{k_{P,R}\pi_P + k_{R^*P,R^*}\pi_{R^*P}}$$

The distribution of duration time at LTR-off states can be expressed:

$$P_{off}(\tau) = k_{R,P}Q(LTR,\tau) + k_{R^*,R^*P}Q(LTR^*,\tau)$$

### 2.13 Parameter values

Table S2-2. Parameter values of LTR-4-state model for experiments testing synergy on HIV reactivation in Figure 4-6, Figure S4-9

| Parameter | Value | | | | |
|---|---|---|---|---|---|
| | Untreated | AC | NE | AC+NE | AC+NS |
| $\gamma$ | 1 | $2.5 \times 10^9$ | 1 | $2.5 \times 10^9$ | $2.5 \times 10^9$ |
| $\alpha$ | 0 | 0 | 1 | 1 | -1 |
| $\omega$ | 100 | | | | |
| $\beta$ | $\beta = 0$ for Detailed Balance; $\beta = 10$ for non-Detailed-Balance | | | | |
| $k_{act}$ | $1 \times 10^{-11}$ | | | | |
| $k_{bindp}$ | 0.001 | | | | |
| $k_{unbindp}$ | 0.01 | | | | |
| $k_{unact}$ | 0.01 | | | | |

Table S2-3. Parameter values of Tat Positive Feedback Loop in Figure 4-6, Figure S4,7,9

| Parameter | Value |
|---|---|
| $k_{mbasal}$ | 0.01 |
| $k_{trs1}$ | 5 |

| | |
|---|---|
| $k_{trs2}$ | 1 |
| $K_{threshold}$ | 75 |
| $\delta$ | 0.01 |
| $k_{unbindp0}$ | 0.01 |
| $d_m$ | 1 |
| $k_{Tat}$ | 10 |
| $d_{Tat}$ | 0.125 |
| Threshold | 75 |
| Total cell number | 10000 |
| Time steps | 100000 |
| Observation time | 100 |

Table S2-4. Parameter values of GFP expression without feedback in Figure 6D-E, Figure S5A-D, S8E-F

| Parameter | Value |
|---|---|
| $k_m$ | 1 |
| $d_m$ | 1 |
| $k_{GFP}$ | 10 |
| $d_{GFP}$ | 0.125 |

### 3. Supplemental Figures

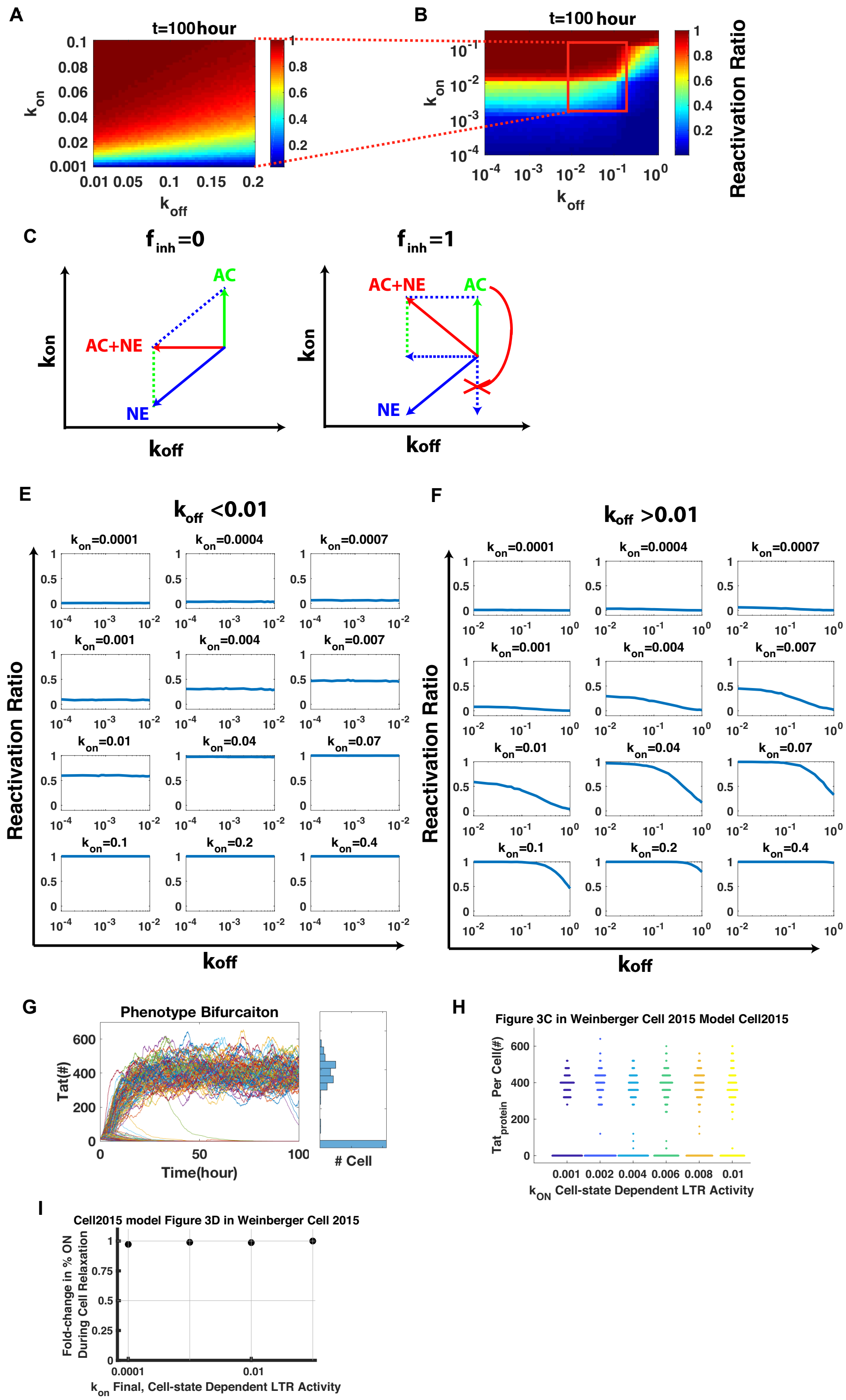

**Figure S1. Refined mesh of the parameter space of synergy on Reactivation of LTR-2-state model as well as verification for the applicability of the model, Related to Figure 2.**

(A-B) Refined heat map of reactivation ratio across different values of $k_{on}$ and $k_{off}$.

(C-D) Vectors in $k_{on}$-$k_{off}$ parameter space indicating adding AC and/or NE with $f_{inh} = 0$ (C) and $f_{inh} = 1$ (D).

(E-F) Reactivation does not change with $k_{off}$ decreasing when $k_{off} < 0.01$; but only increases with $k_{off}$ decreasing when $k_{off} > 0.01$.

(G-I) The LTR-2-state model can explain bimodal distribution of phenotype bifurcation (14) (G) as well as Tat positive feedback being sufficient to control HIV on state despite cellular relaxation from activation to resting (2) (H-I).

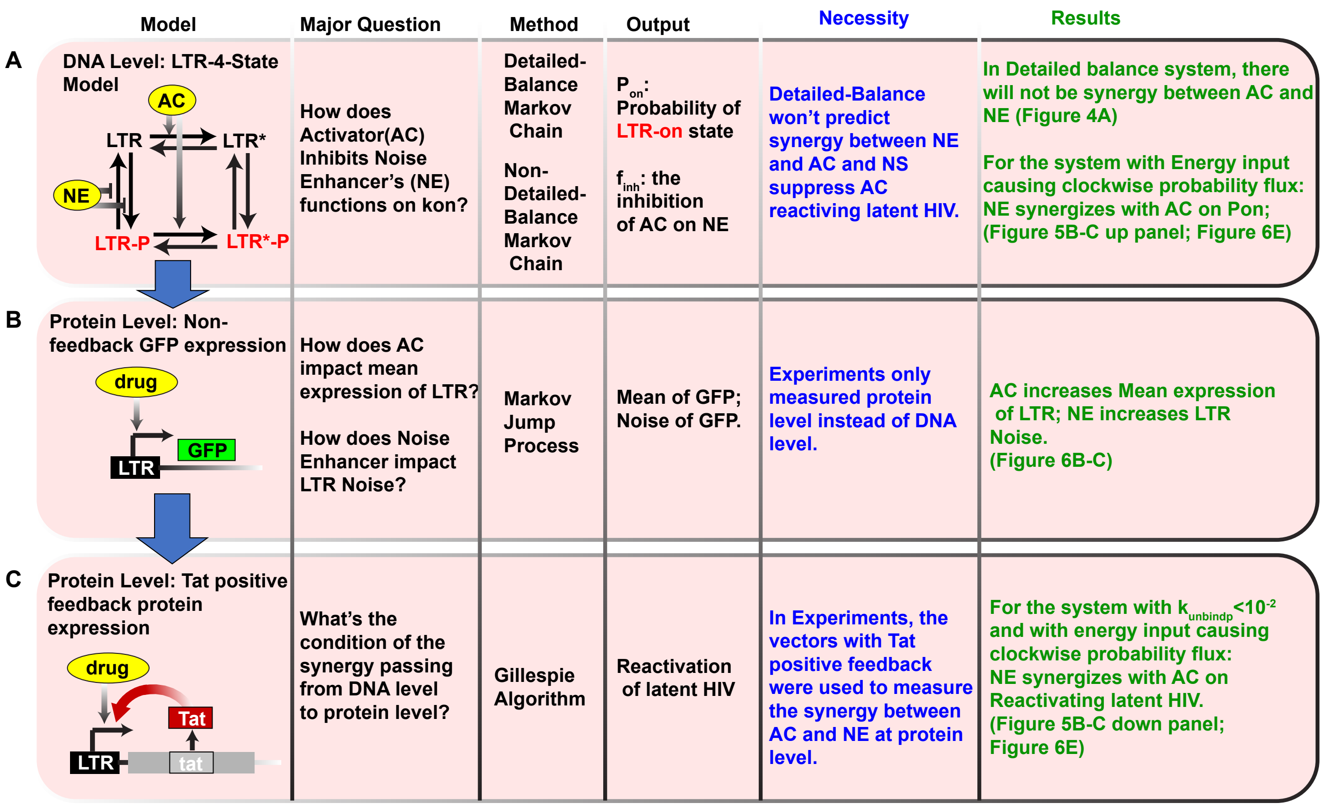

**Figure S2. Progression of LTR-4-state models, Related to Figure 3.**

The flowchart summarizes the two levels of models developed in the main text (Figure 3). (A) is the LTR-4-state model at DNA(gene) level. (B-C) The protein level models without Tat feedback (B) or with Tat feedback (C) are developed, since experiments observed protein expression instead of gene states (1). Collectively, the models predict the synergy between AC and NE in a non-Detailed-Balance model with energy input causing clockwise cyclic probability flux at DNA level and protein level.

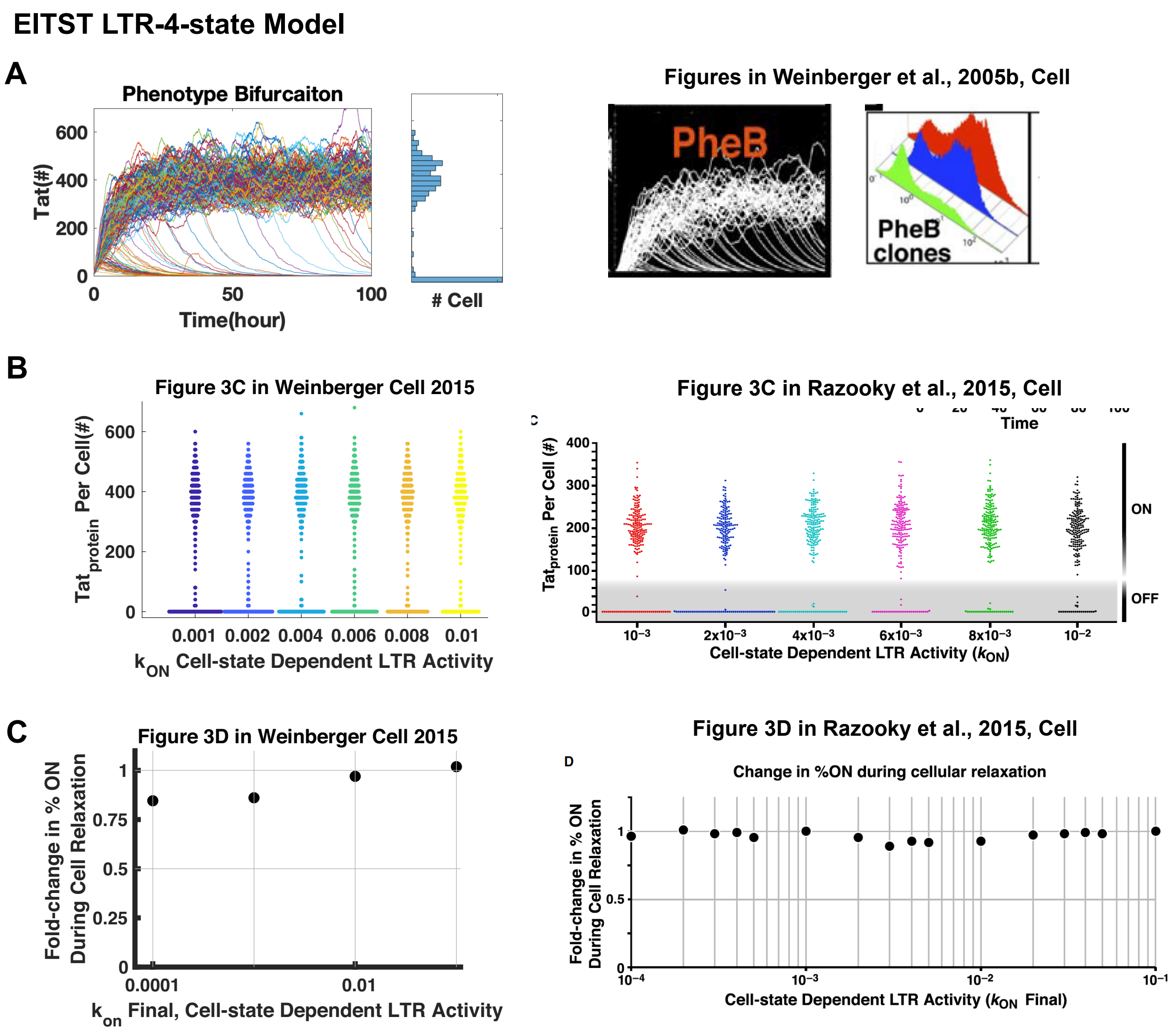

**Figure S3. LTR-4-state model coupled with Tat positive feedback can explain the important previous experiments, related to Figure 3.**

(A-C) The LTR-4-state model (EITST model) can explain bimodal distribution of phenotype bifurcation (14) (A) as well as Tat positive feedback being sufficient to control HIV on state despite cellular relaxation from activation to resting (2) (B-C). On left are our model simulation results and On the right are the previous model results explaining the important experiments.

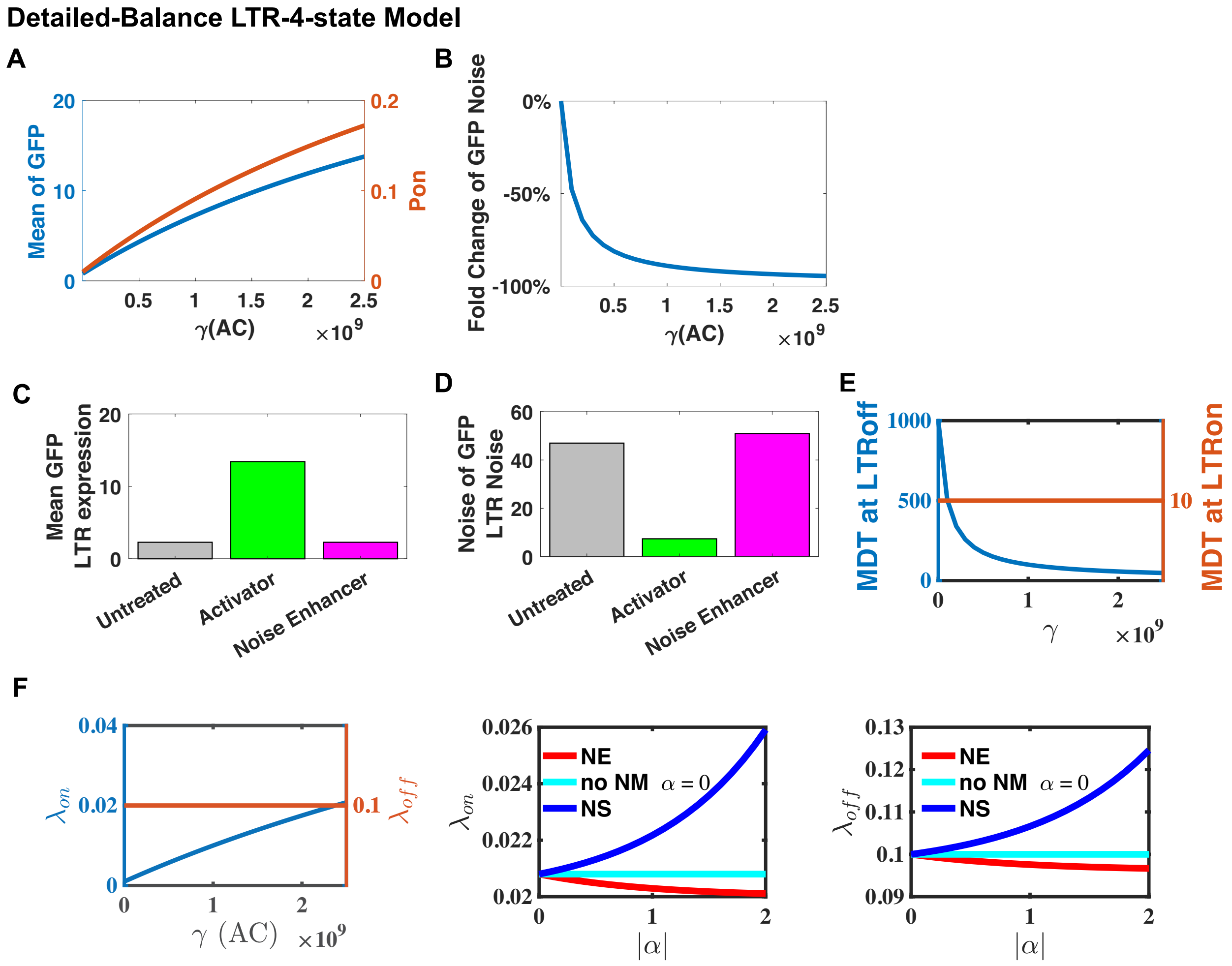

**Figure S4. Mean and Noise of LTR and MDT of LTR-on/off states in Detailed balance LTR-4-state model.**

(A-B) Mean of GFP and Pon (A) as well as the fold change of Noise of GFP (B) changes with $\gamma > 1$ indicating adding AC.

(C-D) Histogram of Mean and Noise of LTR expression for untreated group, group with AC added and group with NE added.

(E-F) MDT at off/on states (E) and the equivalent LTR turning on(or off) rate $\lambda_{on}$(or $\lambda_{off}$) changes with $\gamma > 1$ indicating adding AC. AC mainly increases $\lambda_{on}$.

(G-H) $\lambda_{on}$ (G) and $\lambda_{off}$ (H) changes with $\alpha$ for NE or NS with AC added. NE reduces $\lambda_{on}$ and $\lambda_{off}$ simultaneously while NS increases $\lambda_{on}$ and $\lambda_{off}$ simultaneously.

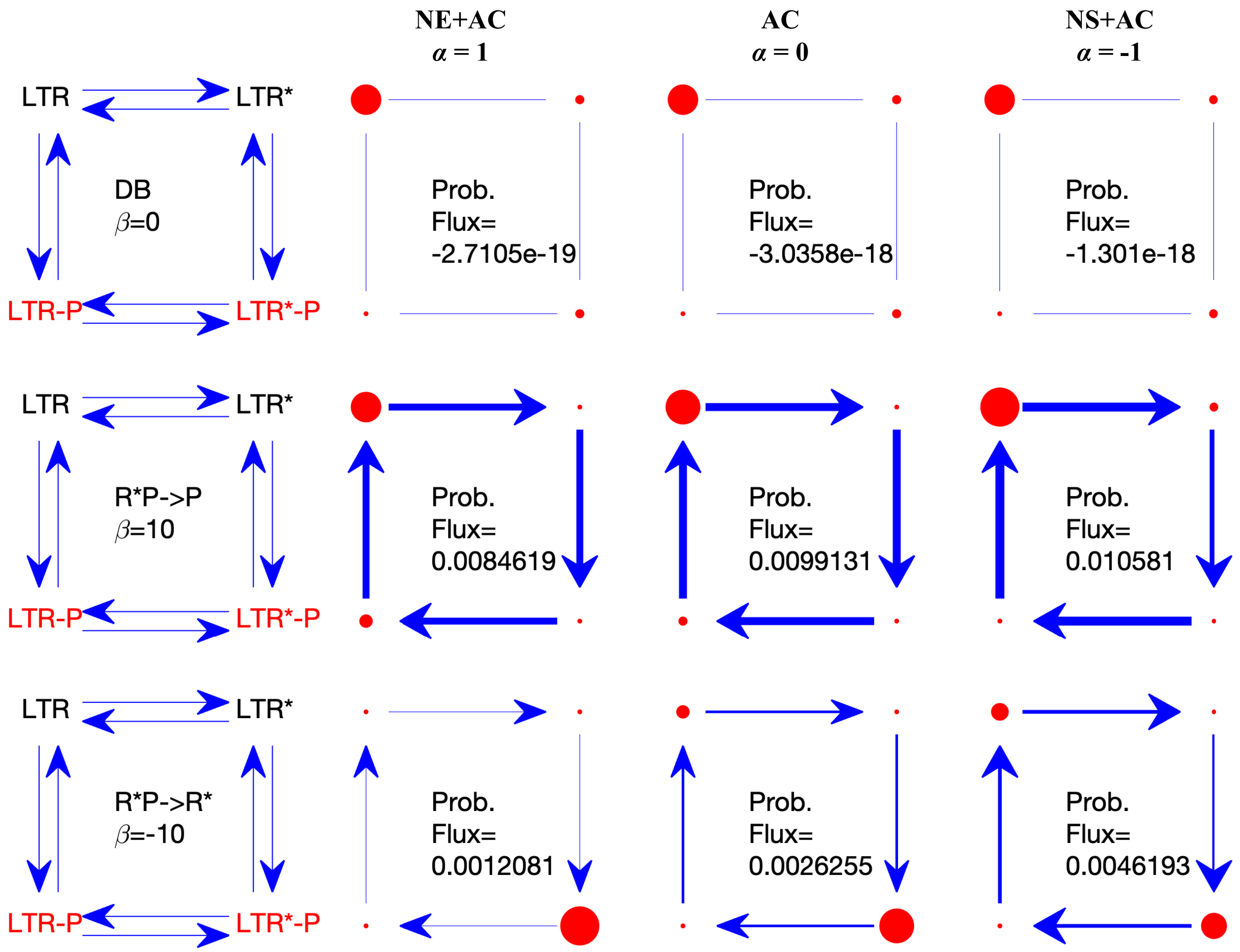

**Figure S5. Non-Detailed-Balance models with clockwise cyclic probability flux, the cycle flux is weakened by NE.**

The first left column represent different non-detailed balance models with clockwise cyclic probability flux. The first line of the text in each model indicates the rate changed by the energy input by multiplying $e^{\beta}$ with the corresponding $\beta$ showed in the second line. The right three columns from left to right represent adding NE and AC ($\alpha = 1$), adding AC ($\alpha = 0$) and adding NS and AC ($\alpha = -1$), respectively. The size of red filled circle indicates the probability of the corresponding state. The width of the arrow indicates the cycle flux.

From left to right, the cycle flux increases with $\alpha$ decreasing.

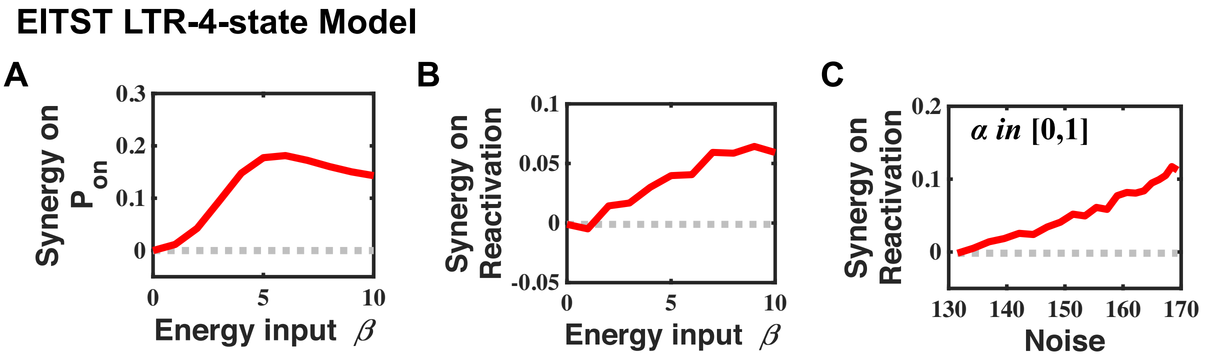

**Figure S6. Synergy on $P_{on}$ and reactivation changes with energy input and Noise of EITST model with even energy distribution.**

(A) The synergy on $P_{on}$ reach the optimal value at $\beta = 10$.

(B) The synergy on Reactivation increases with energy input $\beta$.

(C) The synergy on Reactivation increases with Noise of the system adding only NE with parameter $\alpha \in [0,1]$. These results are calculated from EITST model with $\beta_1 = \beta_2 = \frac{\beta}{2}$.

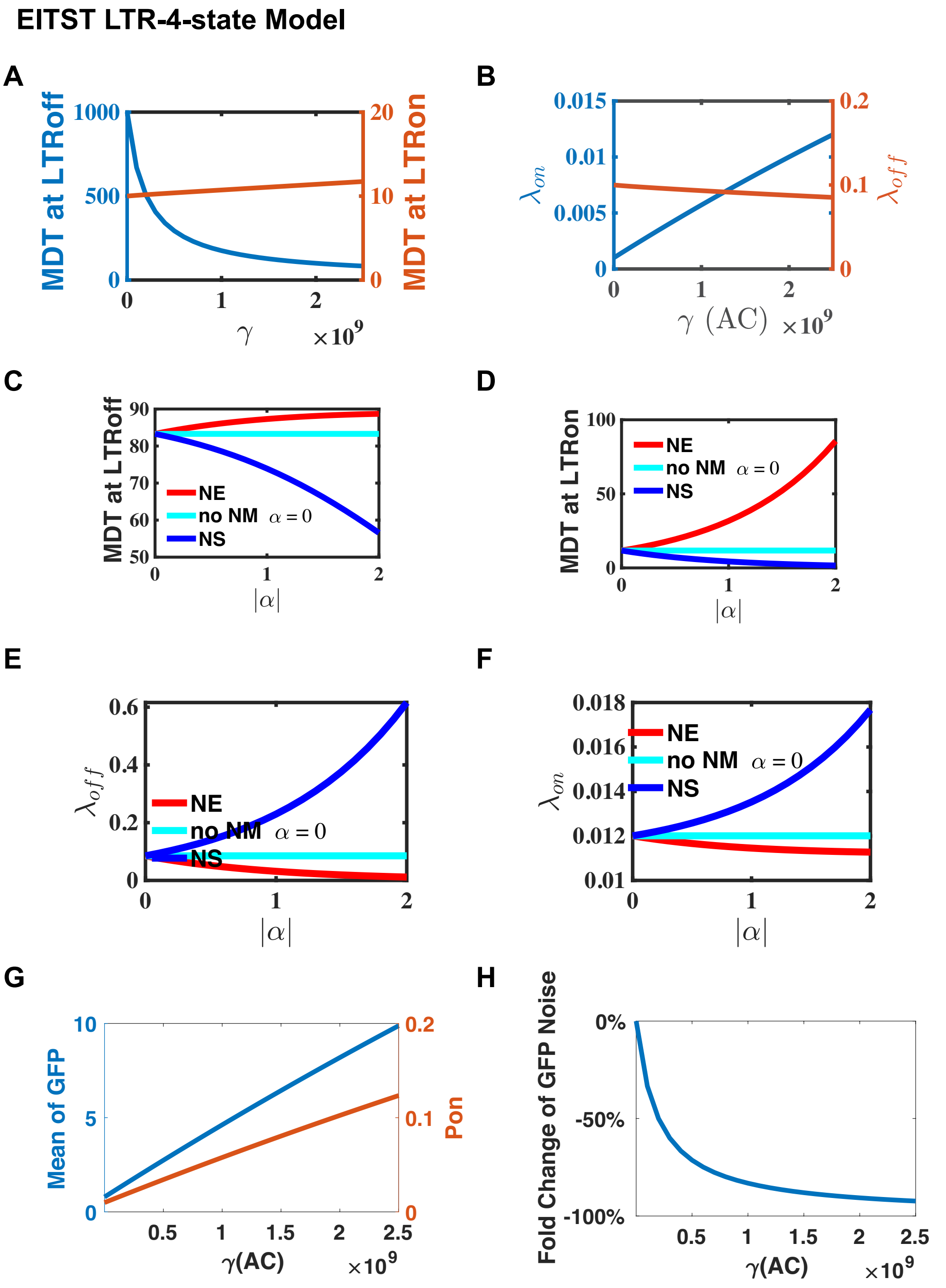

**Figure S7. The MDT at LTR-on/off states of adding AC or NE/NS in EITST LTR-4-state model with even energy distribution.**

(A-B) MDT at off/on states (A) and the equivalent LTR turning on(or off) rate $\lambda_{on}$(or $\lambda_{off}$) changes with $\gamma > 1$ indicating adding AC. AC mainly increases $\lambda_{on}$.

(C-D) Mean Duration Time at LTR-off states and LTR-on states, respectively, of the EITST model.

(E-F) $\lambda_{on}$ (E) and $\lambda_{off}$ (F) changes with $\alpha$ for NE or NS with AC added. NE reduces $\lambda_{off}$ by around 100% and reduces $\lambda_{on}$ by less than 10%, while NS increases $\lambda_{off}$ much more than $\lambda_{on}$.

(G-H) Mean of GFP and Pon (G) as well as the fold change of Noise of GFP (H) changes with $\gamma > 1$ indicating adding AC.

These results are calculated from EITST model with $\beta_1 = \beta_2 = \frac{\beta}{2}$.

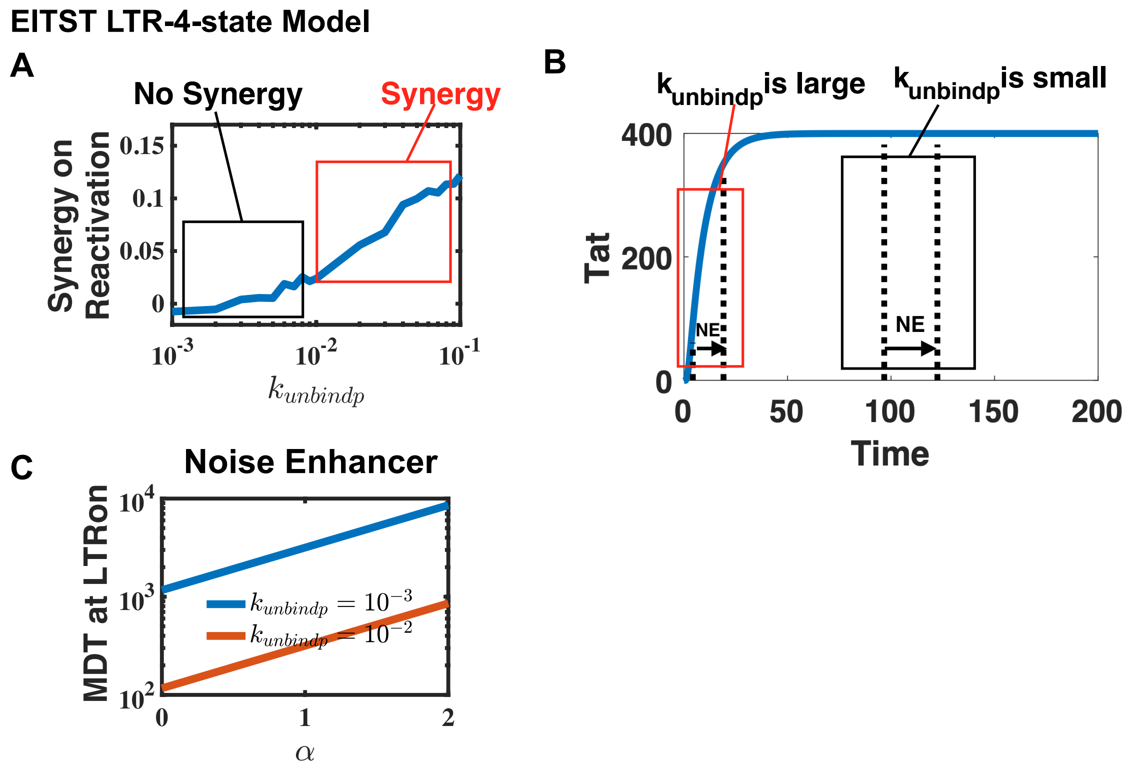

**Figure S8. Synergy only exits when $k_{unbindp} > 10^{-2}$ with even energy distribution.**

(A) Synergy on reactivation is significant (>3%) only when $k_{unbindp} > 10^{-2}$.

(B) At LTR-on states, the mean-field deterministic dynamics of Tat till steady state. It takes Tat 5~25 hours till self-reactivation.

(C) For $k_{unbindp} < 10^{-2}$, the MDT in untreated group ($\alpha = 0$) at LTR-on state is already longer than 100 hours, which is long enough for Tat self-reactivation. Adding NE ($\alpha > 0$) will not increase the benefits.

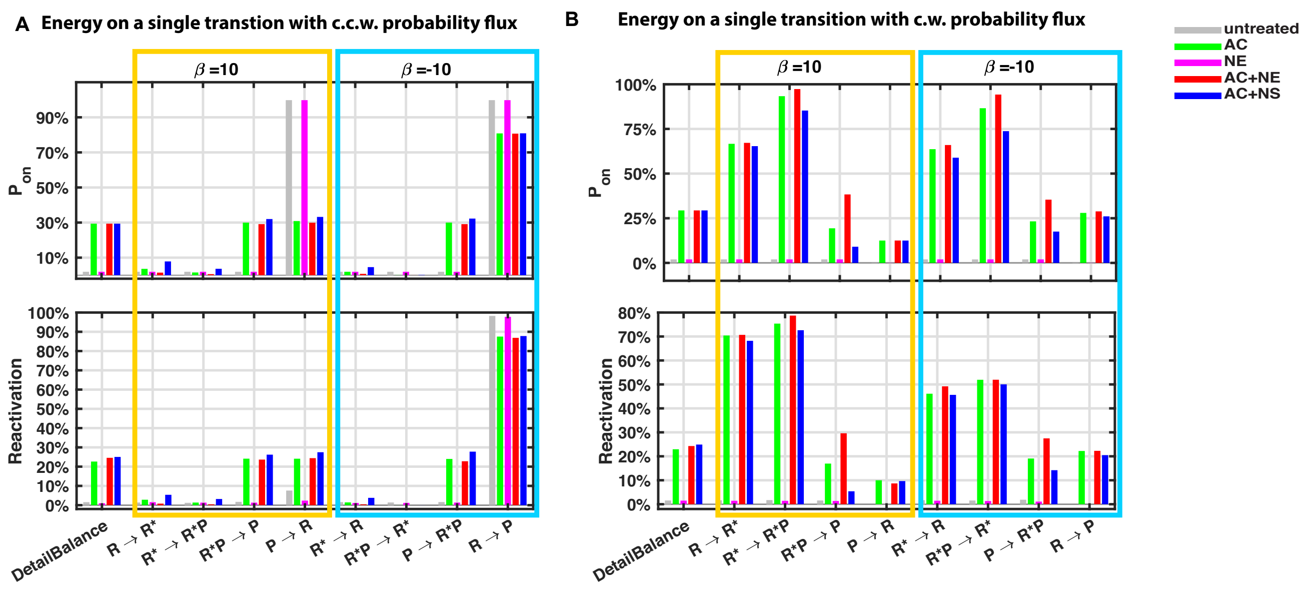

**Figure S9. Tat-binding models predicts the same results: for system with c.w. cyclic probability flux, NE and AC has synergy.**

(A-B) Probability of LTR-on states, $P_{on}$, and Reactivation ratio of Latent HIV, calculated from the Non-Detailed-Balance models coupled with Tat-binding model (See Supplementary Section 2.4 Eqs S2-17) with energy input on different single transitions causing a c.c.w. cycle probability flux or c.w. cycle probability flux, respectively. (See Table S2-1 and Table S1-1 for parameter values.)

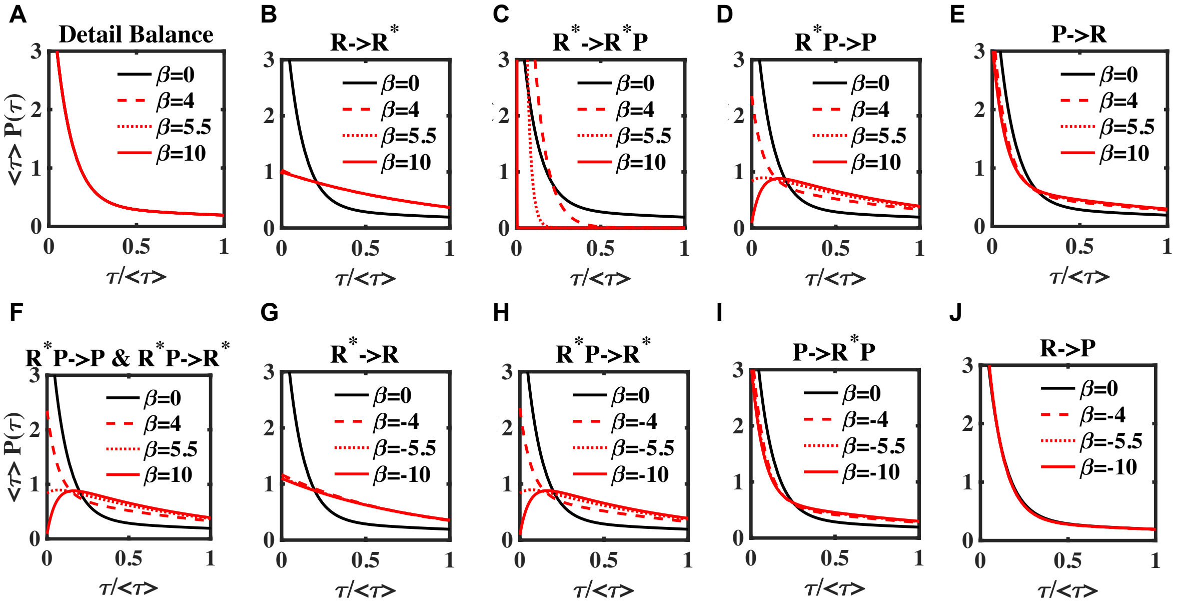

**Figure S10. The behaviors of duration time at LTR-off states distribution $P(\tau)$ for non-equilibrium system models.**

The distribution function starting from being exponential for the equilibrium model (black line) with $\beta = 0$, preserved the monotonicity and convexity for small $\beta$ ($\beta = 4$ for dashed red line). When the energy dissipation $\beta$ increases, the non-equilibrium models can be divided into two categories: (i) (D,F,H) the distribution function $P(\tau)$ shows progressively stronger nonequilibrium signatures as $|\beta|$ increases larger than $\beta_c = 5.3$: concavity and nonmonotonicity appears ($\beta = 5.5$ for dotted red line, $\beta = 10$ for solid red line), (F) is the EITST model with even energy distribution; (ii) (B,D-E,G,I-J) the distribution function $P(\tau)$ still keeps the monotonicity and convexity even though the detailed balanced is broken. (See Table S2-1 and Supplementary Section 2.12 for model details. See Table S2-3 and Table S2-4 for parameter values.)
